# A rubrerythrin locus of *Clostridioides difficile* efficiently detoxifies reactive oxygen species

**DOI:** 10.1101/2024.09.17.613384

**Authors:** Robert Knop, Simon Keweloh, Silvia Dittmann, Daniela Zühlke, Susanne Sievers

## Abstract

As an intestinal human pathogen, *Clostridioides difficile* is the main cause of antibiotic-associated diarrhoea. Endospores of this gram-positive bacterium enter the intestinal tract via faecal-oral transmission, germinate into vegetative and toxin-producing cells and can trigger a *Clostridioides difficile* infection. The microaerophilic conditions (0.1 to 0.4 % O_2_) of the large intestine represent a challenge for the strictly anaerobic organism, which protects itself by a variety of oxidative stress proteins. Four of these are encoded in an operon that is assumed to be involved in the detoxification of H_2_O_2_ and O_2_^●-^. This operon encodes a rubrerythrin (*rbr*), its own transcriptional repressor PerR (*perR*), a desulfoferrodoxin (*rbo*) and a putative glutamate dehydrogenase (*CD630_08280*) with an N-terminal rubredoxin domain, which is only expressed under high oxidative stress conditions.

In this study, the enzyme activity of Rbr, Rbo and CD630_08280 was tested *in-vitro*. Recombinant proteins were overexpressed in *C. difficile* and purified anaerobically by affinity chromatography.

A H_2_O_2_ reduction potential was demonstrated for Rbr, Rbo and glutamate dehydrogenase. Rbr and glutamate dehydrogenase proved to synergistically detoxify H_2_O_2_ very efficiently. Furthermore, Rbo was verified as a O_2_^●-^ reductase and its activity compared to the superoxide dismutase of *E. coli*.

The investigated gene locus codes for an oxidative stress operon whose members are able to completely neutralize O_2_^●-^ and H_2_O_2_ to water and could thus be vital for *C. difficile* to establish an infection in the host.

## Introduction

*Clostridioides difficile* is a Gram-positive, rod-shaped, spore-forming, anaerobic bacterium, which was first isolated from feces of infants in 1935 (1). *C. difficile* and especially its metabolically dormant endospores are known to be omnipresent in the environment and particularly in health care settings (2). As human pathogen it has become one of the most common causes of hospital-acquired infections worldwide with an alarming lethality of 19 % in severe courses (3). The pathogenicity of vegetative cells is based in particular on the expression of two toxins. Both internalize human intestinal epithelial cells through endocytosis and alternate, as a consequence of their glucosyltransferase activity, the actin cytoskeleton, leading to lysis of intestinal epithelial cells (4) and symptoms ranging from mild diarrhea to life-threatening pseudomembranous colitis (5). Upon mucosal damage, the host immune system reacts with a massive inflammation reaction, resulting in the production of reactive oxygen- and nitrogen species (ROS and RNS) to combat the bacterial infection (oxidative burst). Increased levels of ROS such as hydrogen peroxide (H_2_O_2_), hydroxyl radicals (HO^●^) and superoxide (O_2_^●-^) cause oxidative stress and challenge cells to detoxify the ROS to prevent cellular damage (6). In addition to the oxidative stress caused by the inflammatory reaction, the strictly anaerobic bacterium is already exposed to oxygen when it colonises the intestine. It has been found that a healthy human gastrointestinal tract has a longitudinal decreasing O_2_ gradient (7). Regarded as anoxic milieu, there is up to 4 % O_2_ present in the small intestinal lumen, where the germination of *C. difficile* endospores takes place. Further downstream, in the large intestine, the oxygen concentration decreases to 0.1 to 0.4 % allowing for vegetative growth and reproduction of *C. difficile* (7). Considered as strictly anaerobic bacterium, *C. difficile* vegetative cells are known to be sensitive to low O_2_ (8). However, Giordano *et al*. demonstrated a surprisingly high aerotolerance of *C. difficile* 630, as it shows significant growth in a modified hypoxic chamber with up to 2 % O_2_ (9). Other previous studies report that a substantial number of vegetative *C. difficile* cells survive a challenge with approximately 21 % O_2_ for a short period (10, 11). For a long time, it was assumed that the intolerance and sensitivity of anaerobic microorganisms to O_2_ was the consequence of low abundance or complete lack of detoxifying enzymes such as catalase or superoxide dismutase (12). However, such enzymes or the genes encoding them have been found in many members of the anaerobic community (13). The comparably high aerotolerance of *C. difficile* also suggests the presence of an effective oxidative stress response, with an arsenal of genes encoding proteins for the detoxification of several ROS (14). Thus, the human pathogen contains a *sodA* gene that encodes a classical bacterial superoxide dismutase, which has either a manganese or iron centre, depending on metal availability, and catalyses the dismutation of O_2_^●-^ into H_2_O_2_ and O_2_ (15). Typically, hydrogen peroxide is converted to water and O_2_ by catalases, of which *C. difficile* expresses three (*cotCB*; *cotG*; *cotD*), interestingly only during sporulation (16, 17). Both detoxification reactions generate molecular oxygen, which in turn is a potential source for the formation of new ROS. It is therefore not surprising that oxygen-sensitive bacteria use alternative enzymatic pathways for the detoxification of ROS. The bacterial key proteins in this respect are superoxide reductases (SOR) and rubrerythrins, which can gradually transfer electrons from NAD(P)H to ROS via electron carrier protein domains (rubredoxins) and thus reduce O_2_^●-^ (by SOR) to H_2_O_2_ and further to water and NAD^+^ (by rubrerythrins) (Fig.S1) (18–20). Two rubrerythrins (*CD630_08250* (*Rbr*); *CD630_28450* (*Rbr1*) and two reverse rubrerythrins (*CD630_14740*; *CD630_15240*) are encoded in the genome of *C. difficile* 630 (14). One of these rubrerythrins, *rbr*, is the first in a tricistronic operon together with the transcriptional regulator *perR* and the desulfoferrodoxin *rbo* (21), (Fig. 1). This operon is regulated by the peroxide sensing transcriptional repressor PerR, which belongs to the family of iron uptake regulators (Fur) and blocks transcription by binding as a dimer to the DNA binding site. H_2_O_2_ stress causes metal-catalysed histidine oxidation, a conformational change and consequently derepression of the operon (22). The repressor leaves the DNA and the expression of ROS protection proteins takes place (10).

**Figure 1:**
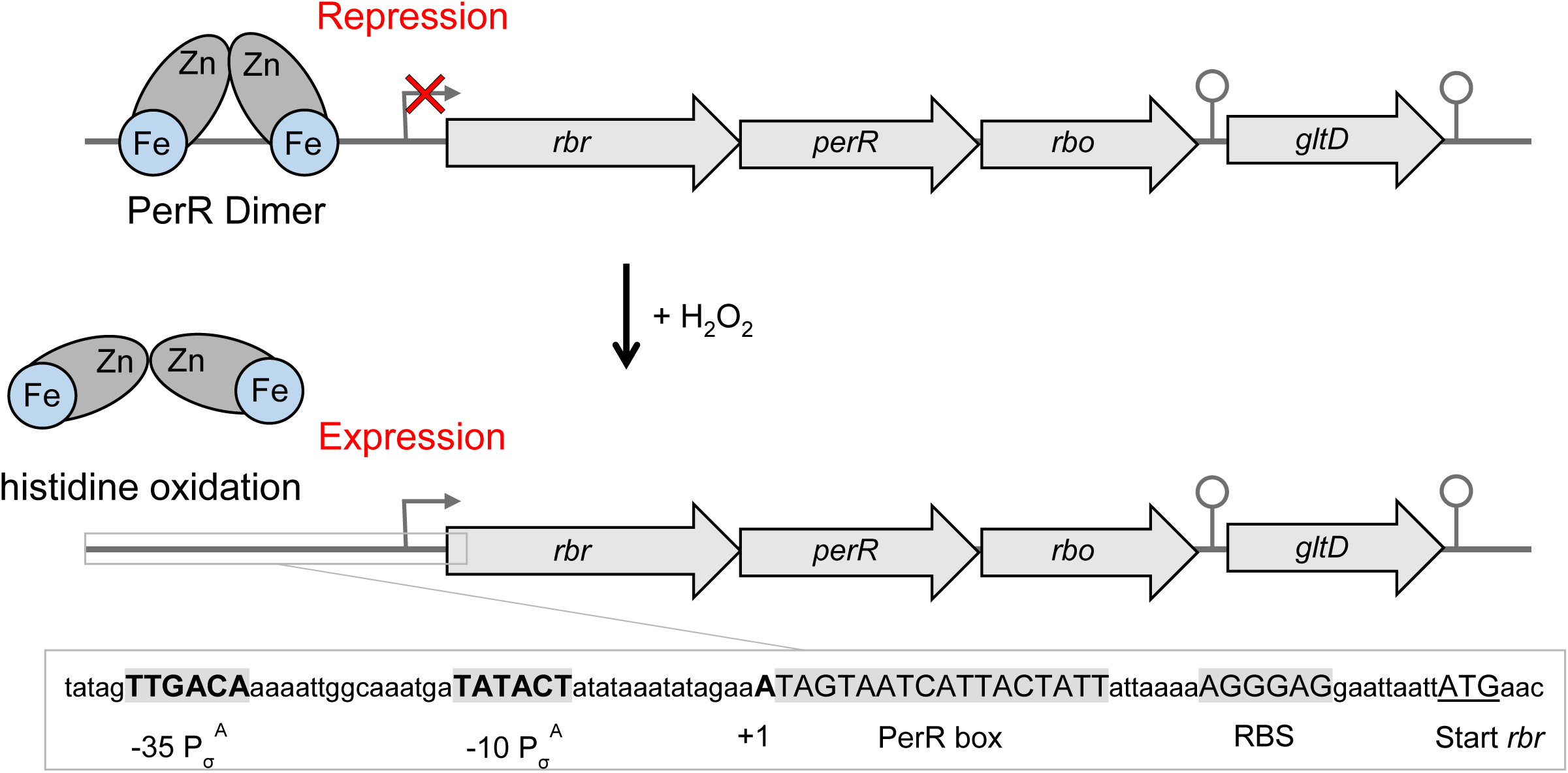
DNA binding of PerR and operon structure with σ^A^ promoter sequence. Mode of PerR function shown schematically. DNA-bound PerR represses the expression of the *rbr* operon. H_2_O_2_ treatment leads to oxidation of PerR, a conformational change and release of the DNA promoter, resulting in expression of the *rbr* operon (adapted from Troitzsch *et al*. (10)). The promoter region is shown below in detail. The -10 and -35 promoter elements recognized by *σ*^A^, a putative PerR binding box and a possible ribosome binding site (RBS) are highlighted in grey. The transcriptional start (gray arrow) and terminator sites (circle) were added as annotated by Soutourina *et al.* and Fuchs *et al*. (21, 24).

The gene *rbo* codes for a desulfoferrodoxin, which were previously called rubredoxin oxidoreductases and were shown to complement superoxide dismutase (SOD) deletion in *Escherichia coli* (25). NAD(P)H-dependent superoxide reductase activity gives desulfoferrodoxins the ability to reduce superoxide to hydrogen peroxide with electrons provided by NAD(P)H (26). A putative SOR activity of the *C. difficile* Rbo may contribute to the surprisingly high aerotolerance of *C. difficile*. Interestingly, downstream of the *rbo* locus an oxidative stress glutamate dehydrogenase with N-terminal rubredoxin fold (*CD630_08280*) is encoded, which is abbreviated as *gltD* in the following. Troitzsch *et al.* were able to show that under strong expression conditions, there is a readthrough of the operon and it is transcribed as a tetracistronic unit together with *gltD* (10). A homologue in *C. acetobutylicum* has already been shown to play an important role in the enzymatic detoxification of ROS (27).

In this work, putative ROS detoxifying proteins of the *perR* regulated operon will be investigated for their activity as stand-alone proteins and in combination with other proteins of the operon *in-vitro*. Specifically, the assumed peroxide neutralizing effect of Rbr and SOR activity of Rbo are to be detected. Furthermore, the role of the oxidative stress glutamate synthase GltD and its possible involvement in the regeneration of Rbr and Rbo will be investigated. The *rbr-perR-rbo-gltD* gene locus possibly constitutes an extensive ROS protection centre being vital for the survival of *C. difficile* in the host.

## Results

### *In-silico* analyses of the *rbr* operon

A number of genes potentially involved in the oxidative stress response of *C. difficile* are located at a specific genetic locus (+1000307-1003256) comprising a rubrerythrin (*rbr*), the peroxide sensing repressor PerR (*perR*), a desulfoferrodoxin (*rbo*) and a N-terminal rubredoxin carrying glutamate synthase (*gltD*). A genome-wide transcriptional start site (TSS) analysis identified a canonical -10 and -35 sequence of a σ^A^-dependent promoter upstream of this operon (Fig. 1) (21). The putative binding sequence for the PerR repressor (PerR box) is located directly between the σ^A^ promoter motif and a possible ribosomal binding site (23). Interestingly, experiments deleting the general stress sigma factor B show *C. difficile* to become more susceptible to ROS, indicating a possible indirect contribution of σ^B^ in the regulation of the operon (28). Troitzsch *et al.* were able to show a readthrough beyond the *rbo* gene in H_2_O_2_-stressed cells (10). This readthrough ensures that the downstream located glutamate dehydrogenase (*gltD*) is expressed only at high stress conditions, which is supported by the fact that no promoter sequence could be identified upstream of *gltD* and that the transcription termination site (TTS) past *rbo* is with -19.10 kcal/mol significantly weaker than the folding energy of *gltD* TTS with -25.20 kcal/mol (21). Using blast analysis (29), we searched for homologs of the four proteins in related clostridia (*C. acetobutylicum; C. beijerinckii; C. botulinum; C. perfringens; C. tetani*) under consideration of their genetic localization and arrangement. The genetic neighborhood consideration revealed that in none of the other clostridia a comparable ROS detoxification operon structure is encoded. For the tested clostridia species, homologous proteins of the *rbr* operon could be found with a percentage identity between 30 and 77 %, except for *C. perfringens* showing no significant similarity to *C. difficile* 630 Rbo (Table 1). Interestingly, a glutamate dehydrogenase with a rubredoxin domain was only found for *C. difficile*. In the other species, only either a large glutamate synthase subunit or, as in the case of *C. perfringens* and *C. tetani*, only the rubredoxin domain were identified as stand-alone proteins with significant similarity. However, a putative oxidative stress glutamate dehydrogenase as a “hybrid” protein could be found in other species than *C. difficile* with high percent identity (65 – 77 %) (Table S1).

**Table 1:** Most similar blast hits of *C. difficile* Rbr; PerR; Rbo and GltD homologs in related clostridia species (status 20^th^ of June 2024).

| Organism | Protein homolog | Protein description | Accession number | protein length [aa] | % identity to *C. difficile* 630 | % similar to *C. difficile* 630 |
| --- | --- | --- | --- | --- | --- | --- |
| *C. difficile* | Rbr | Rubrererythrin | CD630_08250 | 178 | 100 | 100 |
|  | PerR | Peroxide-responsive repressor | CD630_08260 | 129 | 100 | 100 |
|  | Rbo | Desulfoferrodoxin | CD630_08270 | 128 | 100 | 100 |
|  | GltD | Oxidative stress glutamate dehydrogenase | CD630_08280 | 480 | 100 | 100 |
| *C. acetobutylicum* | YotD | Rubrererythrin | KHD38679.1 | 195 | 51 | 68 |
|  | Fur | Fur family transcriptional regulator | WP_185666595.1 | 151 | 30 | 57 |
|  | SodL | Desulfoferrodoxin, superoxide reductase-like | KHD38575.1 | 125 | 41 | 56 |
|  | GltB | Glutamate synthase large subunit | KHD37970.1 | 1507 | 33 | 47 |
| *C. beijerinckii* | YotD | Rubrererythrin | WP_026886601.1 | 195 | 51 | 66 |
|  | Fur | Fur family transcriptional regulator | WP_023974247.1 | 135 | 37 | 56 |
|  | SorL | Desulfoferrodoxin, superoxide reductase-like | AWK52318.1 | 129 | 47 | 64 |
|  | GltB | Glutamate synthase large subunit | AWK49714.1 | 1525 | 34 | 49 |
| *C. botulinum* | Rbr | Rubrererythrin family protein | NFT06462.1 | 179 | 77 | 89 |
|  | Fur | Fur family transcriptional regulator | WP_039258604.1 | 139 | 34 | 55 |
|  | SorL | Desulfoferrodoxin, superoxide reductase-like | WP_106899550.1 | 124 | 44 | 63 |
|  | GltB | Glutamate synthase large subunit | WP_265859470.1 | 1523 | 33 | 49 |
| *C. perfringens* | Rbr | Rubrererythrin | WP_126964111.1 | 179 | 69 | 83 |
|  | PerR | Transcriptional repressor | NWJ12637.1 | 148 | 34 | 55 |
|  | Rbo | ----- No significant similarity found ----- |  |  |  |  |
|  | NorV | Rubredoxin | WP_208373388.1 | 53 | 35 | 50 |
| *C. tetani* | YotD | Rubrererythrin | WP_129009976.1 | 195 | 52 | 71 |
|  | Fur | Fur family transcriptional regulator | WP_023439353.1 | 140 | 34 | 58 |
|  | SorL | Desulfoferrodoxin, superoxide reductase-like | WP_035111706.1 | 124 | 48 | 65 |
|  | NorV | Rubredoxin | WP_115605814.1 | 53 | 37 | 54 |

In all these homologues, the rubredoxin domain is also localized at the N-terminus. Thus, the hybrid protein variant is not exclusively a *C. difficile* feature. All 20 species have a facultative or an even obligate anaerobic lifestyle, indicating a putative evolution from an anaerobic common ancestor.

### Structural analyses of predicted protein fold of Rbr, Rbo and GltD

Rubrerythrins are non-heme iron proteins and known as peroxide stress protectants in a variety of organisms (30). They are built of two iron-binding domains. Firstly, there is a ferritin-like domain consisting of two antiparallel helix pairs connected via an oxo-bridged diiron center, giving the ability to catalyze the oxidation of Fe^2+^ to Fe^3+^. Secondly, there is a rubredoxin like domain that binds another iron atom tetrahedrally via four cysteine residues [Fe(SCys)_4_]. This domain is also involved in electron transfer processes (30). Structural analyses of *C. difficile* Rbr reveals that it counts to the canonical rubrerythrins, having an N-terminal diiron center and a C-terminal rubredoxin domain (Fig. 2A). The position of these two domains can also be switched in so called reverse rubrerythrins (revRbrs), like it is annotated for CD630_14740 and CD630_15240 in *C. difficile* 630.

**Figure 2:**
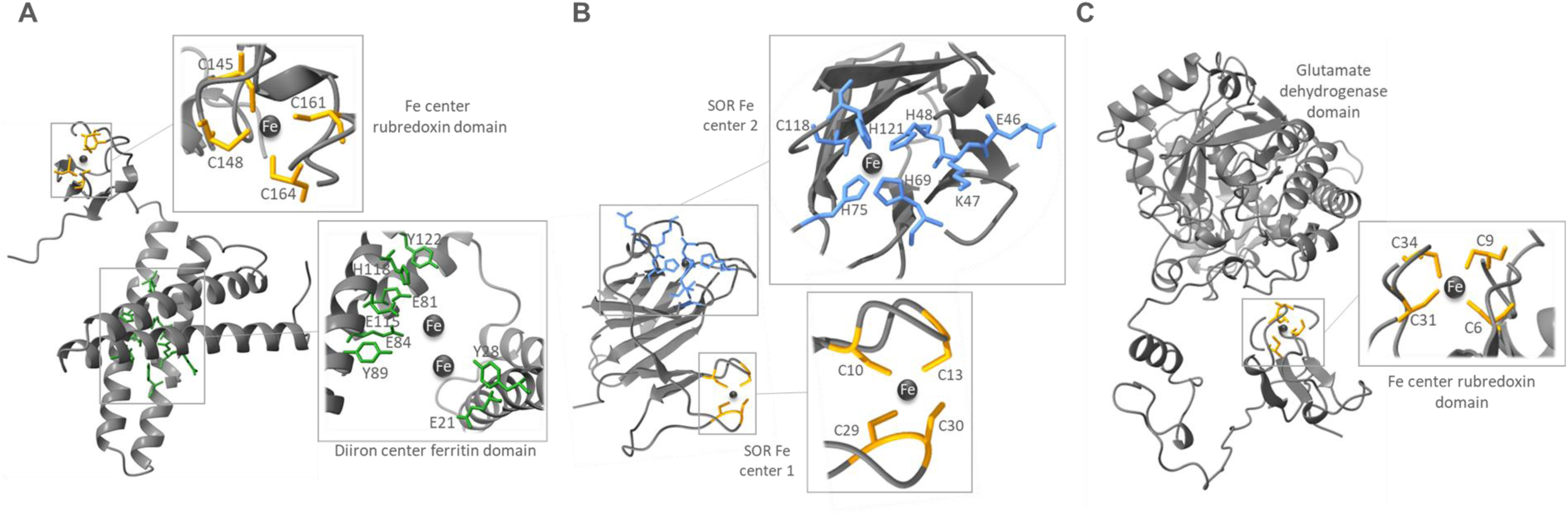
ColabFold predicted protein structures of Rbr (**A**); Rbo (**B**); GltD (**C**) pictured with ChimeraX. Amino acid residues involved in the putative Fe-Ion binding are colored in yellow (rubredoxin domain), green (diiron center ferritin domain) and blue (SOR Fe center 2). Fe-Ions are shown as dark grey circles. SOR: superoxide reductase.

A possible pathway to detoxify highly reactive O_2_^●-^ is via superoxide reductase (SOR) activity of desulfoferrodoxins (Rbo), which were first described in sulphate-reducing bacteria (25, 31). SORs scavenge O_2_^●-^ by reducing it to peroxide taking electrons from NAD(P)H. Regarding the structure and according to the number of two iron centres, *C. difficile* Rbo belongs to the 2Fe-SOR subclass of non-heme iron enzymes (Fig. 2B) (32). The difference to 1Fe-SOR is a [Fe(SCys)_4_] rubredoxin-type centre additional to the mononuclear iron active site [Fe(NHis)_4_(SCys)] (33). The SOR Fe centre 2, whose iron is bound by four histidine and one cysteine residue, is essential for enzymatic activity, whereas the function of the additional SOR Fe centre 1 is not fully understood, but is dispensable for detoxification (34, 35). An amino acid alignment of Rbo to homologous desulfoferrodoxin sequences has shown that several other clostridia as *C. beijerinckii*, *C. tetani* and *C. botulinum* also feature a 2Fe-SOR homolog, but that the enzyme of *C. acetobutylicum* has only the [Fe(NHis)_4_(SCys)] iron site and thus belongs to the 1Fe-SOR, matching the results of Riebe *et al*. (Fig. S2) (27).

Eventually, Figure 2C shows the predicted protein structure of the putative oxidative stress glutamate dehydrogenase GltD of *C. difficile* 630, built of the N-terminal rubredoxin domain with its typical [Fe(SCys)_4_] centre and the C-terminal glutamate dehydrogenase site (Fig. 2C). The physiological role of glutamate dehydrogenase is the reversible NAD(P)^+^ dependent deamination of glutamate to α-ketoglutarate, ammonia and NAD(P)H. One could assume that the NAD(P)H produced by glutamate oxidation could directly be used by the rubredoxin domain of GltD for the recovery of the active iron center of Rbo and Rbr.

### Overexpression and purification of the recombinant proteins of *C. difficile*

To test the enzyme activity and possible concerted action of the proteins to eliminate ROS *in-vitro*, purified protein extracts of Rbr, Rbo and GltD were necessary. Therefore, target proteins were tagged C-terminally with a Twin-Strep-tag^®^ and cloned into the overexpression shuttle vector pDSW1728 under the control of an anhydrotetracycline (ATC) inducible promoter. For an efficient translation, the ribosomal binding site (RBS) of the *slpA* gene of *C. difficile* 630 was inserted in front of the overexpressed recombinant gene sequences. All sequences were verified by sanger sequencing. In order to maintain a protein structure as native as possible and to prevent oxidation of iron centers, recombinant proteins were anaerobically overexpressed in *C. difficile* 630 and purified under anaerobic conditions via affinity chromatography. Figure 3 shows an SDS gel image of the pure recombinant protein fractions (Fig. 3). The identity of proteins was assured by mass spectrometry analysis. The Twin-Strep-tag^®^ has a mass of about 2.8 kDa and causes a small shift in size of recombinant proteins (Rbo: 14.1 kDa; Rbr: 20.6 kDa; CD630_28250: 53.2 kDa). A faint band is visible at 34 kDa in the lane of Rbo, which could be a Rbo dimer as dimerization of desulfoferrodoxin was shown before in *Desulfovibrio desulfurians* (26).

**Figure 3:**
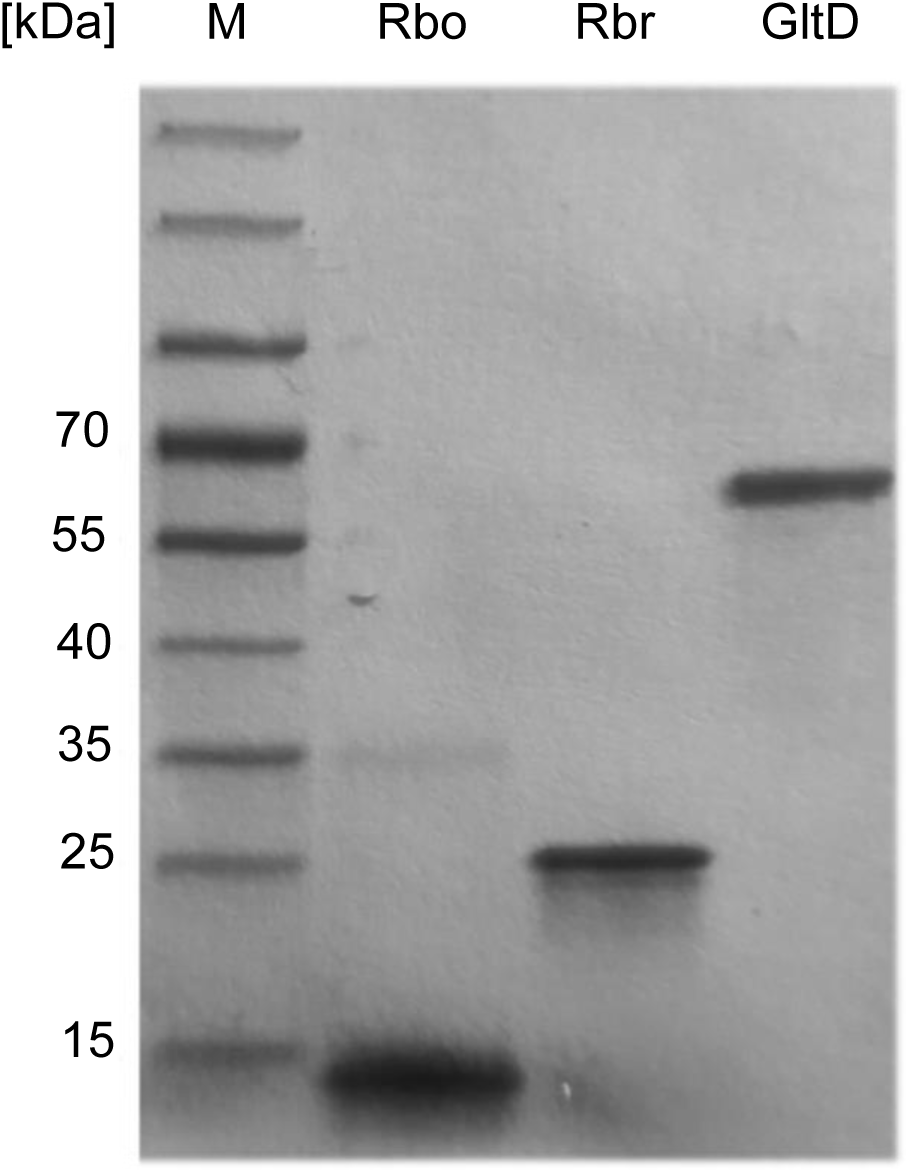
SDS gel image of purified recombinant Rbo, Rbr and GltD of *C. difficile* 630. Proteins were overexpressed from an inducible plasmid in *C. difficile* 630 and purified by affinity chromatography. Pure proteins were visualized by Coomassie brilliant blue staining after separation via SDS-PAGE and identified by mass spectrometry. Theoretical molecular weights: Rbo ∼ 14.1 kDa; Rbr ∼ 20.6 kDa; GltD ∼ 53.2 kDa.

### H_2_O_2_ detoxification activity of Rbr and Rbo

Considering the known ROS detoxification pathway (Fig. S1), we constructed a putative corresponding scheme for *C. difficile*, including all gene products of the *rbr* operon except for regulator PerR (Fig. 4). O_2_^●-^ is reduced to H_2_O_2_ by the superoxide reductase activity of desulfoferrodoxin Rbo. NAD(P)H transfers electrons to Rbo and restores the oxidized catalytic iron center. GltD is possibly the direct donor of NAD(P)H through glutamate oxidation. The degradation pathway of H_2_O_2_ is ensured by rubrerythrin Rbr, which could also reduce H_2_O_2_ generated by Rbo to water. The required electrons could again be donated from NAD(P)H synthesized by GltD.

**Figure 4:**
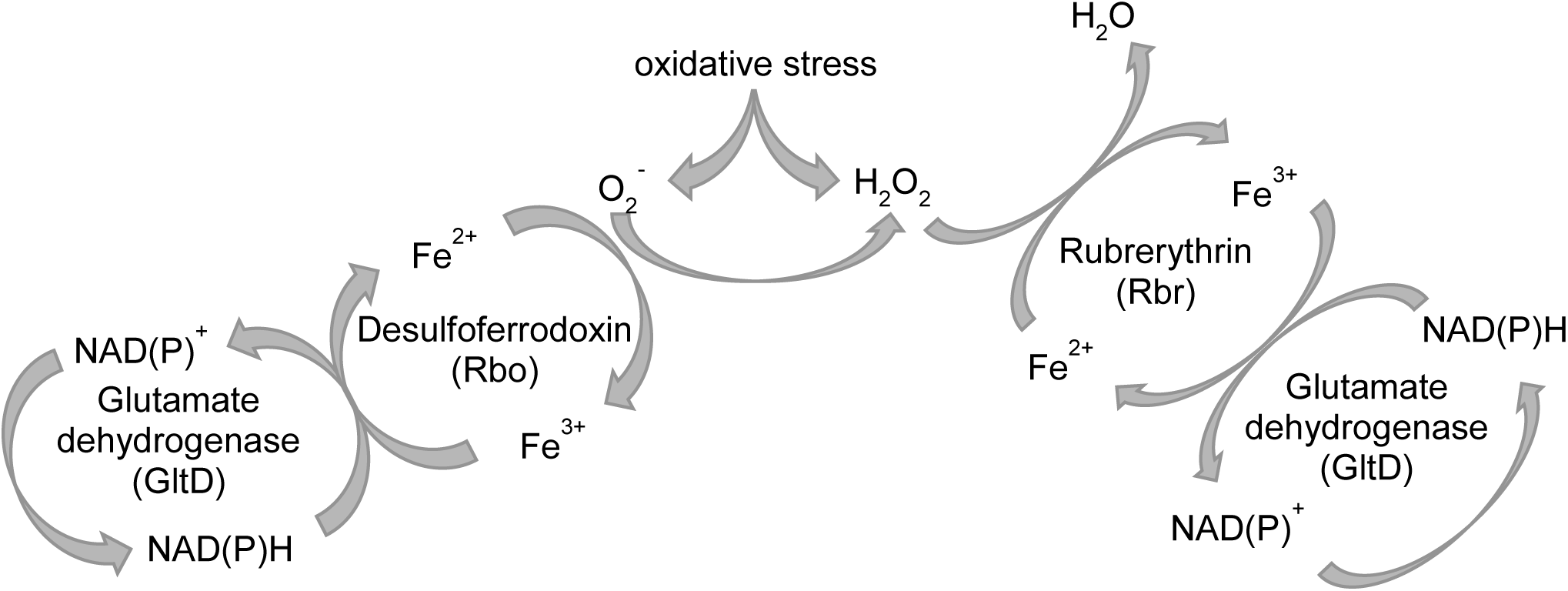
Schematic representation of the putative enzymatic detoxification pathway of H_2_O_2_ and O_2_^-^ by gene products of the *rbr* operon of *C. difficile* (adapted from Troitzsch *et al*. (36)).

The recombinant enzymes were analyzed in defined amounts and in different combinations with each other. All assays were performed in an anaerobic working station. H_2_O_2_ detoxification ability was determined by measuring the consumption of NADH on the basis of the decrease in absorbance at 340 nm (37). H_2_O_2_ was added in a concentration of 100 µM H_2_O_2._ The spontaneous and enzyme-independent reduction of H_2_O_2_ by NADH was considered in a control experiment without any proteins. The blank values determined were subsequently subtracted from all samples in order to separate this basal consumption from actual enzyme activities. After addition of each compound, 10 measurements were taken to see a constant NADH level, starting with NADH, followed by the proteins. H_2_O_2_ was added at last. Furthermore, an experimental run was carried out with 1 μM BSA to make sure NADH is not consumed by any arbitrary protein but due to the specific activity of the recombinant proteins. In addition, the enzymes were analyzed separately in a concentration of 1 µM, to determine their individual effect on NADH concentration. The NADH concentration was normalized to 200 μM to have a uniform reaction starting point, which enables a better comparability of the gathered results. The whole data set of the H_2_O_2_ assays are provided supplementary (Fig. S3). In all experiments, NADH decrease started not before the addition of H_2_O_2_, which demonstrates the applicability of the experimental set-up.

Figure 5 summarizes the results of the H_2_O_2_ detoxification assays of either Rbr or Rbo in combination with GltD in average of three replicates subtracted by the blank values. Rbr and GltD combined, have an effective NADH consumption that clearly exceeds the sum of the values of the single proteins (Fig. 5A). An NADH decrease of 2.9 µM for Rbr, 1.8 µM for GltD and 8.7 µM for the combination of both proteins was measured after 250 sec, whereas the mathematical sum of the Rbr and GltD value is calculated with 4.7 µM.

**Figure. 5:**
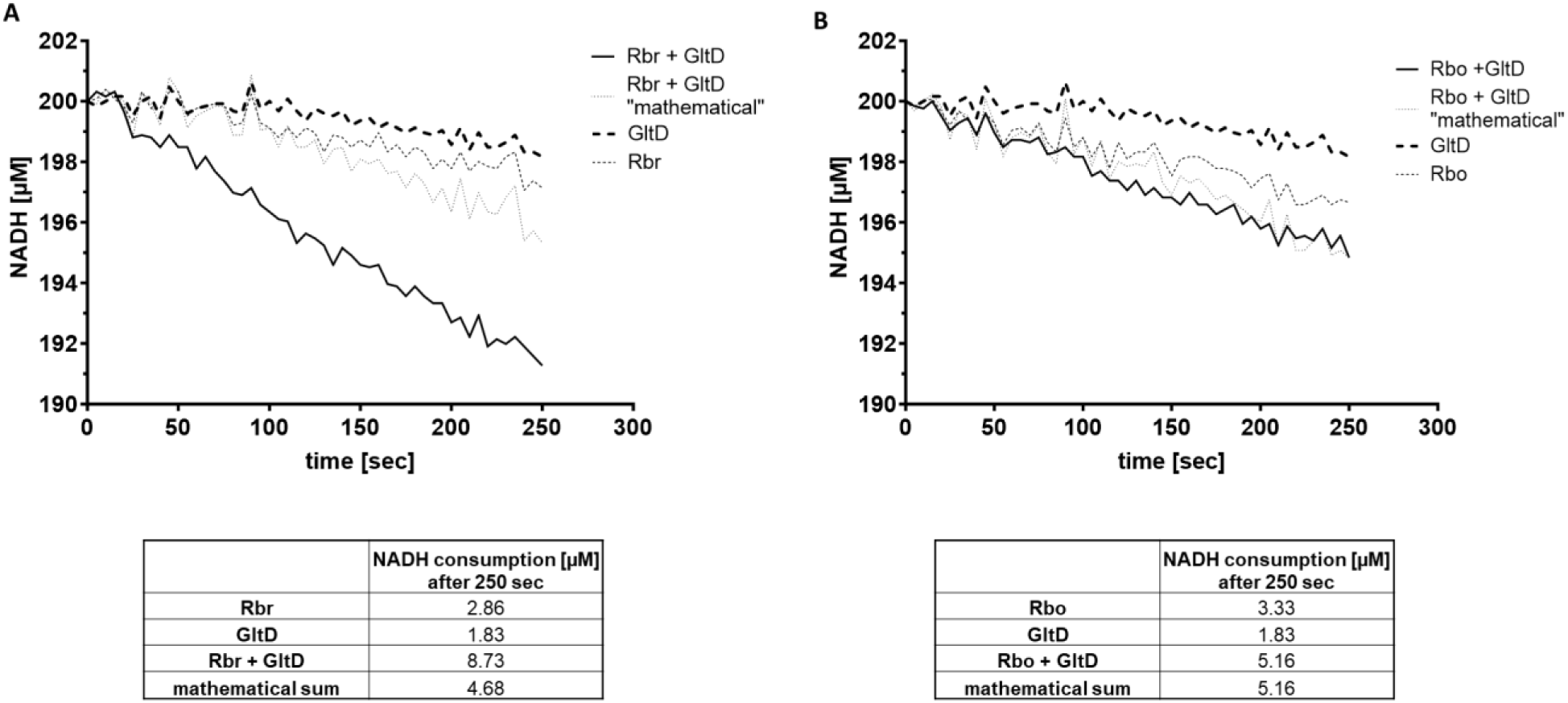
H_2_O_2_ reductase activity of Rbr, Rbo, GltD. The activities were determined anaerobically by measuring NADH consumption monitored at 340 nm every 5 seconds. The graphs presented are the averages from three technical assays for each enzyme. Shown are Rbr, Rbo and GltD individually, their mathematical sum, as well as the measured values of the combination of the two proteins (**A**: Rbr + GltD; **B**: Rbo + GltD). The protein concentrations in the assays were 1 µM for each enzyme (Rbr, Rbo, GltD). Experiments were performed in the presence of 200 µM NADH and 100 µM H_2_O_2_ for the oxidative stress condition. All products were added and the reaction starts with the addition of H_2_O_2_. The determined blank values are subtracted from all samples and normalized to 200 µM NADH.

Moreover, it was shown that GltD has a minor H_2_O_2_ reduction activity and Rbo can also detoxify this ROS even more efficiently than Rbr on its own. However, Rbo and GltD do not seem to cooperate in the context of H_2_O_2_ detoxification, because their joined NADH oxidation rate corresponds to the sum of single rates (Fig. 5B).

### Superoxide detoxification activity of Rbo

To test for the superoxide reduction potential of Rbo of *C. difficile*, we performed an additional assay inducing superoxide stress by the addition of xanthine and xanthine oxidase. O_2_^●-^ is more reactive and not as stable as H_2_O_2_ and has to be generated ad hoc. Xanthine oxidase is able to synthesize large amounts of superoxide by catalyzing the oxidation of xanthine to uric acid through the reduction of O_2_ to O_2_^●-^ (38). Because Xanthine oxidase also reduces NAD^+^ to NADH when oxidizing xanthine, a decreasing concentration of NADH is not a suitable to test for Rbo activity. Hence, another biochemical assay was employed which is the cytochrome c dependent O_2_^●-^ detoxification assay. The assay makes use of the ability of O_2_^●-^ to reduce ferricytochrome c (Fe^3+^) leading to an increased absorbance at 550 nm (Fig. 6) (39).

**Figure 6:**
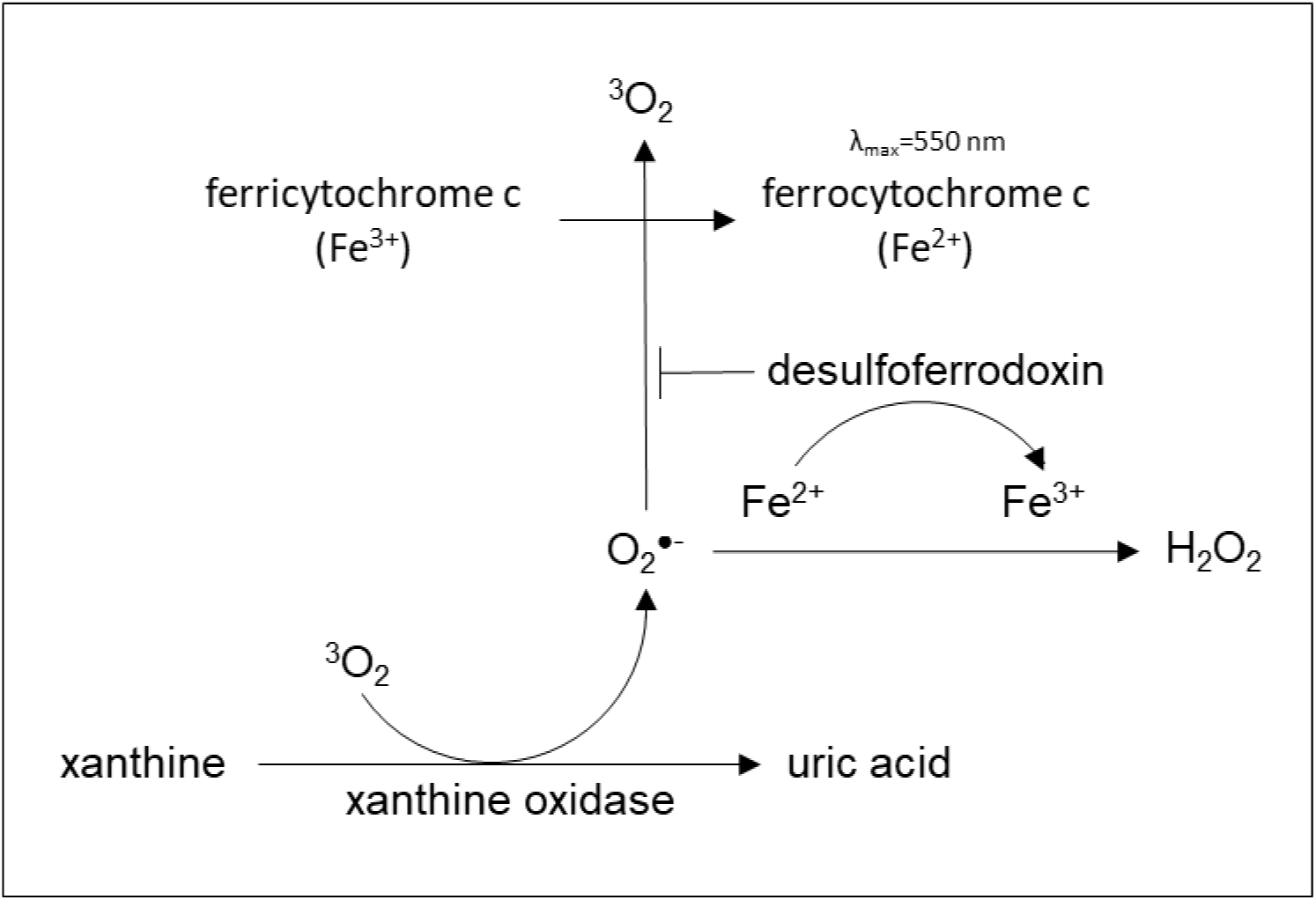
Schematic representation of the superoxide detoxification assay: Xanthine is transformed to uric acid by xanthine oxidase, generating the O_2_^●-^, which in turn reduces ferricytochrome c (Fe^3+^) to ferrocytochrome c (Fe^2+^). The conversion of cytochrome c can be measured at 550 nm (39). Desulfoferrodoxin Rbo scavenges the O_2_^●-^ and ferricytochrome c (Fe^3+^) cannot be reduced. Rbo is oxidized and H_2_O_2_ is produced.

Thus, a reduced absorbance at 550 nm means reduced levels of superoxide. Figure 7 shows the mean of three conducted assays. As expected, in the control sample without Rbo, a significant increase in adsorption at 550 nm was observed immediately after the addition of xanthine oxidase causing superoxide production and cytochrome reduction. The signal at 550 nm increased almost by 60 % in 150 sec after reaction start. In contrast, the presence of 1 µM Rbo causes not only a constant value, but even a slight reduction of 550 nm adsorption. This observation clearly shows that Rbo of *C. difficile* reduces the superoxide anion radical very rapidly impeding the reduction of ferricytochrome c (Fe^3+^) to ferrocytochrome c (Fe^2+^) and does not require the presence of GltD. The slight decrease in the colorimetric curve after the start of the reaction is most likely caused by H_2_O_2_ produced by Rbo, which oxidizes traces of ferrocytochrome c (Fe^2+^) to ferricytochrome c (Fe^3+^) (40). In order to better evaluate and relate the detoxifying effect of Rbo, the assay was also carried out with superoxide dismutase SodB of *E. coli*, which was overexpressed and also purified by affinity chromatography using a C-terminal located Twin-Strep-tag^®^ and is known to efficiently detoxify O_2_^●-^. As expected, the addition of SodB can also prevent the reduction of cytochrome c reflected in the missing rise of absorbance at 550 nm. SodB is a superoxide dismutase that does not produce any H_2_O_2_. Thus, there is no slight drop in the curve after xanthine oxidase addition as it was observed for Rbo. The results of the individual replicates are summarised in the supplement (Fig. S4).

**Figure 7:**
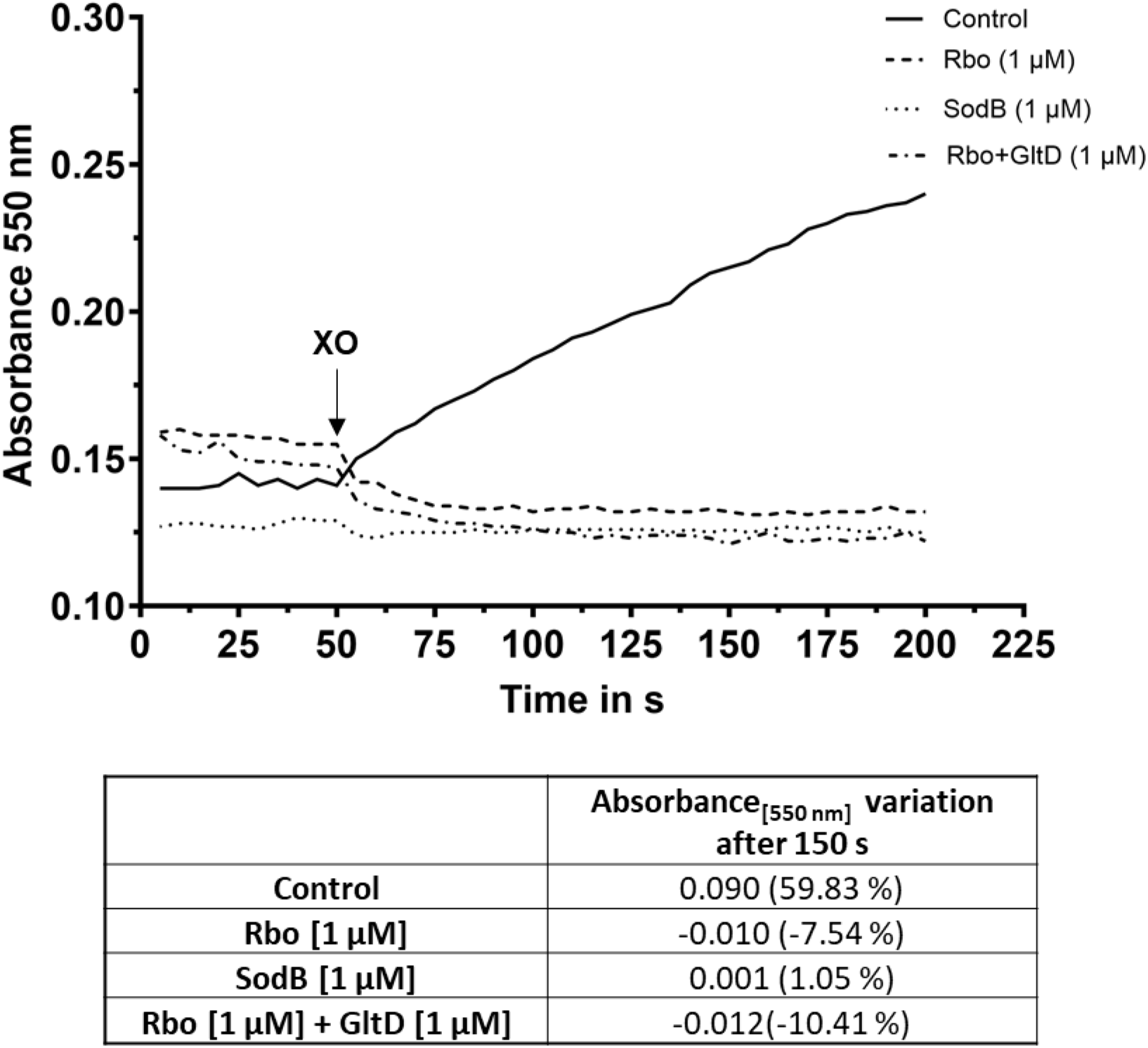
Superoxide reductase activity of Rbo and SodB measured via absorbance at 550 nm indicating cytochrome c reduction. Shown are Rbo and SodB individually, as well as the measured values of the combination of Rbo + GltD. The protein concentrations in the assays were 1 µM for each enzyme (Rbo, GltD, SodB). Experiments were performed in the presence of 200 µM NADH, 500 µM xanthine, 20 µM cytochrome c and 15 µg/ml xanthine oxidase (XO) added to start the generation of superoxide. The graphs represent the average of three replicates. The absorbance variation over 150 s after reaction start is shown in the table below.

Since both Rbo and SodB could eliminate all nascent O_2_^●-^ at once in concentrations of 1 µM, smaller amounts of the enzymes were added to compare the efficiency of the enzymes of *C. difficile* and *E. coli.* For this purpose, the assays were carried out with four different protein amounts (1 µM; 0.1 µM; 0.01 µM; 0.001 µM) (Fig. 8). Figure 8A shows quite distinctly that 0.1 µM Rbo is still sufficient to reduce all O_2_^●-^ before it can convert cytochrome c. However, 0.01 µM (increase of 31 %) and 0.001 µM (increase of 38 %) of Rbo do not suffice anymore to prevent cytochrome c reduction. The *E. coli* enzyme SodB is not as efficient as Rbo and can scavenge only part of the O_2_^●-^ with an intermediate increase in A_550nm_ at a SodB concentration of 0.1 µM. The results of each replicate are given supplementary (Fig. S5).

**Figure. 8:**
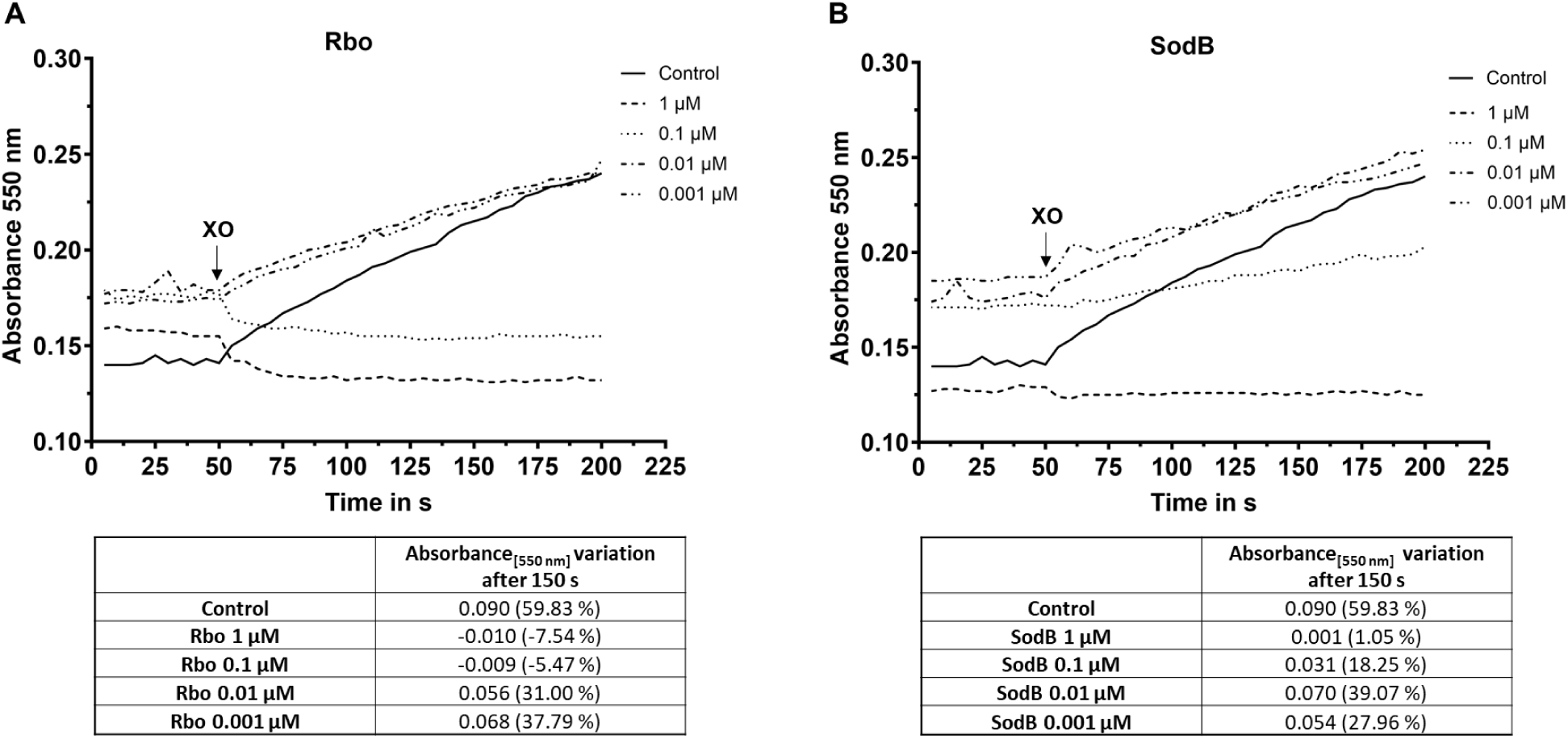
Superoxide reductase activity of Rbo (**A**) and SodB (**B**) measured via absorbance at 550 nm every 5 seconds in a concentration gradient. Protein concentrations of 1 µM; 0.1 µM; 0.01 µM and 0.001 µM were tested. Experiments were performed in the presence of 200 µM NADH, 20 µM cytochrome c, 500 µM xanthine and 15 µg/ml xanthine oxidase (XO) for the superoxide generation. The graphs represent the average of two replicates. The absorbance variation over 150 s after reaction start is given in the tables below.

## Discussion

The anaerobic enteropathogen *C. difficile* is able to tolerate atmospheric oxygen exposure for a surprisingly long time (9). Just like many other anaerobic microorganisms, *C. difficile* has various strategies to actively protect itself against O_2_ and toxic ROS. Certain proteins with their detoxifying effect, of which *C. difficile* has a whole arsenal, play a key role here (23). Two flavodiiron proteins (FdpA, FdpF) and two reverse rubrerythrins (revRbr1, revRbr2) were found, which not only generally play an important role in ROS tolerance, but exhibit different reductase activities depending on the O_2_ concentration (41). Furthermore, three thioredoxin systems have been identified in *C. difficile*, of which two are considered to be involved in the oxidative stress response (42). In the genome of *C. difficile* 630, one locus in particular stands out as it encodes a whole operon of oxidative stress proteins (43). In this study, we focused on the enzymatic members of this specific operon, i.e. a rubrerythrin (Rbr), a desulfoferrodoxin (Rbo) and a glutamate dehydrogenase with an N-terminal rubredoxin fold (GltD) and investigated their potential to detoxify the ROS H_2_O_2_ and O_2_^●-^, thus contributing to the aerotolerance of the pathogen.

### Rbr: the canonical rubrerythrin is part of an operon

In addition to Rbr, *C. difficile* encodes three other rubrerythrins (CD630_14740; CD630_15240; CD630_28450), which are neither arranged in an operon structure nor are they controlled by the repressor PerR (21). Recently, rubrerythrins CD630_14740 and CD630_15240 together with two flavodiiron proteins (FdpA and FdpF) were shown to be induced by O_2_ and putatively repressed by the transcriptional regulator CD630_17770 (OseR) (41). The four rubrerythrins in *C. difficile* are controlled by either *σ*^A^ (*rbr*), by the general stress response sigma factor *σ*^B^ (*CD630_14740*) or by both (*CD630_15240*) (41). The exception is the remaining classical rubrerythrin *CD630_28450*, which is under the control of *σ*^G^. Its expression could so far only be detected in endospores (44). CD630_14740 and CD630_15240 are highly identical reverse rubrerythrins sharing an amino acid sequence identity of 96 %, but are identical by only 33 % to the canonical Rbr. In this work, we were able to show that Rbr alone can effectively reduce H_2_O_2_ and that its efficiency can even be increased by the addition of GltD, regarding the three times higher NADH consumption. This observation indicates that H_2_O_2_ is substrate for Rbr, in accordance with its regulation by PerR, which is oxidized by H_2_O_2_. Whether the varying arrangement of rubredoxin and ferritin domains in reverse and canonical rubrerythrins is decisive for different enzyme activities remains to be investigated. Knockout studies have shown that deletion of both reverse rubrerythrins leads to an increased sensitivity in the presence of low O_2_ in *C. difficile*, indicating a functional redundancy of these two very similar enzymes (14). A lower peroxide tolerance is to be expected in a *rbr* mutant, which will be investigated in future studies. However, none of the genes of the *rbr-perR-rbo-gltD* operon have been predicted to be essential according to a transposon analysis in *C. difficile* strain R20291 (45). This could mean that none of the gene products is needed for basic cellular functions but just in times of oxidative stress or that there are backup systems that take over when the gene products are missing. However, when the *rbr-perR- rbo-gltD* operon is entirely derepressed, as it is the case in strain *C. difficile* 630*Δerm* (10), Rbr is the second most abundant protein after the S-layer protein SlpA (46), indicating a very efficient expression of the operon and its importance in ROS detoxification. The huge amount of Rbr in the H_2_O_2_-stressed cell could compensate for a possibly lower efficiency of the protein to detoxify H_2_O_2_ compared to other rubrerythrins.

### Rbo, the desulfoferrodoxin in *C. difficile*

In the literature, the desulfoferrodoxin of *C. difficile* is known under various names: Dsr for desulfoferrodoxin (21), Sor for superoxide reductase (23,32) or Rbo (rubredoxin oxidoreductase), which is valid in the Uniprot database and was thus used in this work. All these names apply to the same O_2_^●-^ detoxifying protein encoded by gene *CD630_08270*, the only desulfoferrodoxin in *C. difficile*. Homologous desulfoferrodoxins have been identified in a few hundred bacteria and archaea, of which the majority have an anaerobic lifestyle (32).

Biological membranes are not permeable to O_2_^●-^ (47), so it is assumed that the main source of intracellular O_2_^●-^ is of cellular origin (48) when molecular oxygen is reduced by one electron. Then fast detoxification of O_2_^●-^ by Rbo becomes important. Rbo is located in an operon, regulated by a repressor sensing H_2_O_2_ stress. However, for the gram-positive pathogen *Staphylococcus aureus* it has been reported, that PerR also detects O_2_ and facilitates H_2_O_2_ sensing of PerR (49) which could be especially relevant for strictly anaerobic bacteria. Considering this and the fact that O_2_^●-^ detoxification in *C. difficile* yields H_2_O_2_, location of *rbo* in an operon that is regulated by PerR appears plausible.

A very recently published paper by Kochanowsky *et al.* has shown that a *rbo* deletion mutant is more sensitive to O_2_^●-^ and that heterologous overexpression of the enzyme in *E. coli* results in increased tolerance of *E. coli* to O_2_^●-^ (32). In this study, we could overexpress Rbo homologously in *C. difficile* and could isolate the protein in high purity which was the basis to prove O_2_^●-^- reductase activity *in-vitro*. Our biochemical assays could show that Rbo reduces O_2_^●-^ very efficiently in the presence of NADH but that no other cellular component is required. In a direct comparison with O_2_^●-^ detoxifying superoxide dismutase SodB of *E. coli*, Rbo proved to be more efficient which can be explained by the strictly anaerobic lifestyle of *C. difficile* compared to facultative anaerobiosis of *E. coli* and by the different enzymatic reaction mechanisms, since Rbo is an O_2_^●-^ reductase generating H_2_O_2_, that has to be detoxified by rubrerythrins for instance, whereas SodB belongs to the class of dismutases generating O_2_ which is not toxic for *E. coli*.

Rbo was furthermore reported to be induced by cathelicidin LL-37, an antimicrobial peptide of the innate human immune system, suggesting its involvement in the infection process (50). However, proteomic studies of a *rbo* deletion mutant have shown that the deletion of the O_2_^●-^ reductase does not seem to have an effect on sporulation, toxin production or resistance to certain antibiotics, like metronidazole, cefoxitin, and trimethoprim (32).

### GltD is more than a glutamate dehydrogenase

It has been described that a NADH-rubredoxin oxidoreductase (NROR) of *C. acetobutylicum* can efficiently reduce clostridial rubredoxin (51). Therefore, NROR is a key enzyme in the ROS detoxification pathway as it regenerates the rubredoxin domain of desulfoferrodoxins and rubrerythrins by consuming NADH. We postulated that GltD, a glutamate dehydrogenase of *C. difficile* with an N-terminal rubredoxin domain, could act as NROR of rubrerythrin Rbr and desulfoferrodoxin Rbo which are encoded in the same genetic locus. Especially in times of high oxidative stress conditions, NADH availability is scarce. A big advantage of GltD as a NROR would be its ability to directly produce the required NADH by the conversion of L-glutamate to ammonium and *α*-ketoglutarate thereby generating reduction equivalents in form of NADH. The importance for this ROS detoxification pathway is substantiated by the fact that GltD is only expressed under strong oxidative stress conditions by readthrough of a transcriptional terminator in front of the gene. Oxidative stress-induced GltD expression was not only shown by Northern blot analyses (10) but also by proteome data of *C. difficile* 630Δ*erm*, a strain that constitutively expresses the *rbr-perR-rbo-gltD* due to a loss of function in the transcriptional repressor PerR (46).

A blast analysis has shown that some anaerobic species feature a hybrid protein as GltD most likely to deal with oxidative stress and to ensure the NADH supply for the reduction of ROS. However, *C. difficile* also codes for another glutamate dehydrogenase (GluD), which is secreted and routinely used in diagnostics to detect the human pathogen in stool samples using a GDH ELISA (glutamate dehydrogenase enzyme-linked immunosorbent assay) (52). Both proteins are just 21 % identical and the rubredoxin domain is lacking in GluD. Still, a higher H_2_O_2_ sensitivity could be shown in a *gluD* deletion mutant (53). If NADH shortage is reason for the higher H_2_O_2_ sensitivity of the mutant or if GluD is otherwise involved in the oxidative stress response still has to be investigated. Likewise, the effect of a *gltD* deletion and a possibly higher sensitivity of the mutant to ROS is subject to future studies.

In this study, we could show that GltD itself has a small H_2_O_2_ reduction capability. Strikingly, it significantly enhances the potential of Rbr to reduce H_2_O_2_ to an extent much more than the additive effect of both enzymes indicating that GltD could act as NROR for Rbr regeneration. Regarding Rbo, we determined a good potential of the enzyme to reduce O_2_^●-^. An addition of GltD did not lead to an increased activity of Rbo, or the used experimental setup did not enable us to detect it (data not shown). If GltD cannot transfer electrons to Rbo, there are different possible reasons. One could be the Twin-Strep-tag^®^ on the C-terminus that was added to all recombinant proteins for affinity chromatography purification. It might block possible interaction sites. An N-terminal affinity tag or no protein tagging but purification with protein specific antibodies could clarify this issue. However, it is also possible that Rbo requires another NROR than GltD for its reduction. There are further putative NRORs (CD630_1623; CD630_1157) postulated in *C. difficile*, which could be involved in the O_2_^●-^ detoxification pathway by reducing Rbo (27). Eventually, GltD might not require a NROR at all, since it belongs to the 2Fe-SOR class featuring a rubredoxin centre that could directly receive electrons from NADH.

In summary, in this study three enzymes encoded in an operon involved in the oxidative stress response of *C. difficile* could be overexpressed homologously in *C. difficile* and purified by affinity chromatography providing anaerobic conditions at all times. Employing different biochemical assay, we could demonstrate the proposed roles in the detoxification pathways of ROS (Fig. 4) showing that Rbr efficiently reduces H_2_O_2_, that Rbo detoxifies O_2_^●-^ and that GltD significantly enhances Rbr activity, probably by acting as NROR of the rubrerythrin. The *rbr-perR-rbo-gltD* gene locus thus represents an important tool to fight the ROS challenge during infection and complements the arsenal of detoxifying mechanisms that are already known in *C. difficile* (14, 53, 54). An increased expression of *rbr* and *rbo* in mouse and porcine animal models also indicates an important role of the gene products for survival in the host (55, 56). Further studies are required to fully understand the importance of the investigated proteins in pathogenesis. However, an increased oxygen tolerance of pathogenic anaerobes has been widely appreciated as virulence factor (57). Hence, the *rbr-perR-rbo-gltD* operon could be an attractive target site for novel antimicrobial strategies in CDI therapy.

## Material and Methods

### Bioinformatical methods

Detailed *in-silico* analyses of gene sequences in *C. difficile* 630 were performed using the PRODORIC genome browser (58). Multiple amino acid sequence alignments of protein sequences were generated using the Clustal Omega tool (59). The prognosis of transcriptional terminator sites (TTS) was calculated with the “*ARNold Finding Terminators*” web service (60) and compared with a list of TTS annotated by Fuchs *et al*. (21). Prediction of the protein fold was carried out with the ColabFold platform (61) and completed with a domain analysis done by the ScanProsite tool (62). To find homologs in other species we used the blast search tool (National Library of Medicine).

### Bacterial strains and growth conditions

All studies were carried out using strains *C. difficile* 630 (DSM 27543), *E. coli* DH5α (DSM 6897) and ST18 (DSM 22074), which were obtained from the German Collection of Microorganisms and Cell Cultures GmbH (DSMZ) (Braunschweig, Germany).

*C. difficile* 630 was routinely cultured anaerobically in Brain Heart Infusion (BHI) broth or on BHI agar plates (1.7 % agar) inside an anaerobic chamber (95 % N2 and 5 % H2) from Don Whitley Scientific Ltd. (Bingley, England) at 37 °C. If necessary, additional thiamphenicol was added to the medium at a final concentration of 25 μg/ml. *E. coli* cultures were grown aerobically in lysogeny broth (LB) or on LB agar plates (1.7 % agar) supplemented as needed with 25 µg/ml chloramphenicol and 50 µg/ml 5-aminolevulinic acid for the selection in the ST18 strain which was used as donor strain for plasmid conjugation into *C. difficile* 630 (63).

### Plasmid and overexpression strain construction

All plasmids and DNA oligonucleotides used in this study are listed in Table S2 and S3. For the construction of overexpression plasmids target genes (*CD630_08270 (rbr)*; *CD630_08270 (rbo)*; *CD630_08280(gltD)*) were amplified from genomic DNA of *C. difficile* 630 using specific primers. Forward primers included the ribosome binding site (RBS) of *C. difficile* SlpA, and a SacI restriction site. The reverse primer includes a BamHI restriction site and a Twin-Strep-tag^®^ for a latter purification of the overexpressed proteins by a strep-tactin column. The PCR product and the *E. coli*-*C. difficile* shuttle vector pDSW1728 (64) were digested with SacI and BamHI (Thermo Fisher Scientific, Waltham, MA, USA) and fused, resulting in the according tetracycline-inducible overexpression vector (pDSW1728-*rbr*; pDSW1728-*rbo*; pDSW-*gltD*). Sequences were confirmed by Sanger sequencing. The plasmid was transformed into the donor strain *E. coli* ST18 (63) following mating in *C. difficile* 630 to construct the overexpression mutants.

### Expression and purification of recombinant *C. difficile* proteins

8 ml BHI medium supplemented with 0.1 % cysteine and 0.01 % taurocholic acid were inoculated with 200 µl of spore suspension. After 24 h of incubation at 37 °C, fresh BHI was inoculated with different volumes of germinated spores and grown overnight. The following morning, precultures having an optical density (OD_600_) between 0.9 and 1.1 were used to inoculate 400 ml main cultures to an OD_600_ of 0.05. Overexpression of target proteins was induced by the addition of 200 ng/μl anhydrotetracycline (ATC) at an OD_600_ of 0.1. After reaching an OD_600_ of 1.0 samples were harvested, filled into gas-tight centrifuge tubes (TPP, Switzerland) and centrifuged at 8500 rpm at 4 °C for 3 minutes. During the following steps, care was taken to ensure that samples did not come into contact with atmospheric oxygen to prevent an oxidation of iron containing proteins. Cell pellets were resuspended in anaerobic buffer W (100 mM Tris-HCl, 150 mM NaCl; pH 7.5) and mechanical disrupted in three homogenization cycles (each 30 s, 6.5 m/s) using a FastPrep-24 5G (MP Biomedicals, LLC, Irvine, CA, USA). Cell debris and glass beads were removed by centrifugation (13.300 rpm, 10 min, 4 °C). Protein extracts were subjected to affinity chromatography according to the manufacturer’s protocol using the “Strep-Tactin^®^XT High Capacity column” (IBA Lifesciences GmbH; Germany) and eluted in buffer E (100 mM Tris; 150 mM NaCl; 50 mM Biotin; pH 7.5). Protein quantification was performed using the ROTI^®^ Nanoquant system (Carl Roth) which is based on the Bradford method (65). To determine the purity of the protein samples, they were separated according to their size using an SDS-PAGE. The identity of proteins was assured by mass spectrometry analysis (66).

### Biochemical activity assays

To avoid oxidation of iron-containing proteins by atmospheric O_2_, all analyses were carried out strictly anaerobically. For this purpose, of all buffers and solutions O_2_ was removed by nitrogen gassing. The experiments were carried out in an anaerobic workstation at 37 °C. All assays were performed in three replicates if not stated otherwise.

### Spectrophotometric measurement of the H_2_O_2_-reductase activity

H_2_O_2_ reduction activity was tested for Rbr, Rbo and GltD. Since NADH is converted to NAD^+^ during the reaction, protein activity was deduced from the decreasing NADH concentration, i.e. the at 340 nm. In order to continuously determine the NADH concentration, the reactions were carried out in 1 ml glass cuvettes in a spectrophotometer. For a final volume of 1 ml, NADH should be at a concentration of 200 μM, the proteins at 1 μM each and H_2_O_2_ at 100 μM in buffer C. Absorbance was determined in 5 sec intervals, with 10 values measured after the addition of each component. This was to show that the NADH remains stable. After addition of H_2_O_2_, 50 more values were recorded to track the decrease in NADH concentration over time.

### Xanthine oxidase assay for superoxide scavenging activity of Rbo and SOD

The assay was based on a previously published protocol to investigate superoxide dismutase activity (67). Xanthine oxidase catalyzes the oxidation of xanthine to uric acid, reducing oxygen to the superoxide anion radical, which in turn transfers its unpaired electron to cytochrome c. Ferricytochrome (Fe^3+^) is thus reduced to ferrocytochrome c (Fe^2+^). The amount of reduced cytochrome c can be measured photometrically at 550 nm. The assay was performed in a 1 ml plastic cuvette, into which aerobic buffer C (50 mM Tris-HCl; pH 7,5) was introduced and subsequently cytochrome c [20 µM], NADH [200 µM], xanthine [500 µM] and the purified proteins were added at the appropriate concentrations. Absorbance was measured in 5 sec intervals to establish a baseline. Superoxide generation was started with the addition of xanthine oxidase [15 µg/ml) and absorbance was measured every 5 sec for further 150 seconds.

### Statistical analyses

Differences between the protein activity were tested for statistical significance using multiple t testing. α was set to 0.05.

Visualization of statistical analyses was performed using GraphPad Prism software (GraphPad Software Inc., La Jolla, CA)

## Supplemental Material

Supplemental material to this article is available online.

## Supporting information

Knop et al_supplements

## Acknowledgements

This work was supported by the German Research Foundation (453440095 to S.S.). We declare no conflict of interest.

## References

1. Hall IC. INTESTINAL FLORA IN NEW-BORN INFANTS. Am J Dis Child 1935; 49(2):390.

2. Vedantam G, Clark A, Chu M, McQuade R, Mallozzi M, Viswanathan VK. *Clostridium difficile* infection: toxins and non-toxin virulence factors, and their contributions to disease establishment and host response. Gut Microbes 2012; 3(2):121–34.

3. RKI. Infektionsepidemiologisches Jahrbuch meldepflichtiger Krankheiten für 2020.

4. Carter GP, Rood JI, Lyras D. The role of toxin A and toxin B in *Clostridium difficile*-associated disease: Past and present perspectives. Gut Microbes 2010; 1(1):58–64.

5. Smits WK, Lyras D, Lacy DB, Wilcox MH, Kuijper EJ. *Clostridium difficile* infection. Nat Rev Dis Primers 2016; 2:16020.

6. Bardaweel SK, Gul M, Alzweiri M, Ishaqat A, ALSalamat HA, Bashatwah RM. Reactive Oxygen Species: the Dual Role in Physiological and Pathological Conditions of the Human Body. Eurasian J Med 2018; 50(3):193–201.

7. Theriot CM, Young VB. Interactions Between the Gastrointestinal Microbiome and *Clostridium difficile*. Annu Rev Microbiol 2015; 69:445–61.

8. Holý O, Chmelař D. Oxygen tolerance in anaerobic pathogenic bacteria. Folia Microbiol (Praha) 2012; 57(5):443–6.

9. Giordano N, Hastie JL, Carlson PE. Transcriptomic profiling of *Clostridium difficile* grown under microaerophillic conditions. Pathog Dis 2018; 76(2).

10. Troitzsch D, Zhang H, Dittmann S, Düsterhöft D, Möller TA, Michel A-M et al. A Point Mutation in the Transcriptional Repressor PerR Results in a Constitutive Oxidative Stress Response in *Clostridioides difficile 630Δerm*. mSphere 2021; 6(2).

11. Edwards AN, Karim ST, Pascual RA, Jowhar LM, Anderson SE, McBride SM. Chemical and Stress Resistances of *Clostridium difficile* Spores and Vegetative Cells. Front Microbiol 2016; 7:1698.

12. McCord JM, Keele BB, Fridovich I. An enzyme-based theory of obligate anaerobiosis: the physiological function of superoxide dismutase. Proc Natl Acad Sci U S A 1971; 68(5):1024–7.

13. Briukhanov AL, Thauer RK, Netrusov AI. Katalaza i superoksiddismutaza v kletkakh strogo anaérobnykh mikroorganizmov. Mikrobiologiia 2002; 71(3):330–5.

14. Kint N, Alves Feliciano C, Martins MC, Morvan C, Fernandes SF, Folgosa F et al. How the Anaerobic Enteropathogen *Clostridioides difficile* Tolerates Low O2 Tensions. mBio 2020; 11(5).

15. Li W, Wang H, Lei C, Ying T, Tan X. Manganese superoxide dismutase from human pathogen *Clostridium difficile*. Amino Acids 2015; 47(5):987–95.

16. Permpoonpattana P, Phetcharaburanin J, Mikelsone A, Dembek M, Tan S, Brisson M-C et al. Functional characterization of *Clostridium difficile* spore coat proteins. J Bacteriol 2013; 195(7):1492–503.

17. Permpoonpattana P, Tolls EH, Nadem R, Tan S, Brisson A, Cutting SM. Surface layers of *Clostridium difficile* endospores. J Bacteriol 2011; 193(23):6461–70.

18. Hillmann F. Von obligater Anaerobiose zur Aerotoleranz: Die oxidative Stressantwort von *Clostridium acetobutylicum*. neue Ausg. Saarbrücken: Suedwestdeutscher Verlag fuer Hochschulschriften; 2010.

19. Jenney FE, Verhagen MF, Cui X, Adams MW. Anaerobic microbes: oxygen detoxification without superoxide dismutase. Science 1999; 286(5438):306–9.

20. Riebe O, Fischer R-J, Wampler DA, Kurtz DM, Bahl H. Pathway for H2O2 and O2 detoxification in *Clostridium acetobutylicum*. Microbiology (Reading) 2009; 155(Pt 1):16–24.

21. Fuchs M, Lamm-Schmidt V, Sulzer J, Ponath F, Jenniches L, Kirk JA et al. An RNA-centric global view of *Clostridioides difficile* reveals broad activity of Hfq in a clinically important gram-positive bacterium. Proc Natl Acad Sci U S A 2021; 118(25).

22. Lee J-W, Helmann JD. The PerR transcription factor senses H2O2 by metal-catalysed histidine oxidation. Nature 2006; 440(7082):363–7.

23. Kint N, Morvan C, Martin-Verstraete I. Oxygen response and tolerance mechanisms in *Clostridioides difficile*. Curr Opin Microbiol 2022; 65:175–82.

24. Soutourina OA, Monot M, Boudry P, Saujet L, Pichon C, Sismeiro O et al. Genome-wide identification of regulatory RNAs in the human pathogen *Clostridium difficile*. PLoS Genet 2013; 9(5):e1003493.

25. Pianzzola MJ, Soubes M, Touati D. Overproduction of the rbo gene product from Desulfovibrio species suppresses all deleterious effects of lack of superoxide dismutase in *Escherichia coli*. J Bacteriol 1996; 178(23):6736–42.

26. Pinto AF, Rodrigues JV, Teixeira M. Reductive elimination of superoxide: Structure and mechanism of superoxide reductases. Biochim Biophys Acta 2010; 1804(2):285–97.

27. Riebe O, Fischer R-J, Bahl H. Desulfoferrodoxin of *Clostridium acetobutylicum* functions as a superoxide reductase. FEBS Lett 2007; 581(29):5605–10.

28. Kint N, Janoir C, Monot M, Hoys S, Soutourina O, Dupuy B et al. The alternative sigma factor σB plays a crucial role in adaptive strategies of *Clostridium difficile* during gut infection. Environ Microbiol 2017; 19(5):1933–58.

29. Altschul SF, Madden TL, Schäffer AA, Zhang J, Zhang Z, Miller W et al. Gapped BLAST and PSI- BLAST: a new generation of protein database search programs. Nucleic Acids Res 1997; 25(17):3389– 402.

30. deMaré F, Kurtz DM, Nordlund P. The structure of *Desulfovibrio vulgaris* rubrerythrin reveals a unique combination of rubredoxin-like FeS4 and ferritin-like diiron domains. Nat Struct Biol 1996; 3(6):539–46.

31. Romão CV, Liu MY, Le Gall J, Gomes CM, Braga V, Pacheco I et al. The superoxide dismutase activity of desulfoferrodoxin from *Desulfovibrio desulfuricans* ATCC 27774. Eur J Biochem 1999; 261(2):438–43.

32. Kochanowsky R, Carothers K, Roxas BAP, Anwar F, Viswanathan VK, Vedantam G. *Clostridioides difficile* superoxide reductase mitigates oxygen sensitivity. J Bacteriol 2024; 206(7):e0017524.

33. Emerson JP, Cabelli DE, Kurtz DM. An engineered two-iron superoxide reductase lacking the Fe(SCys)4 site retains its catalytic properties in vitro and in vivo. Proc Natl Acad Sci U S A 2003; 100(7):3802–7.

34. Pinto AF, Romão CV, Pinto LC, Huber H, Saraiva LM, Todorovic S et al. Superoxide reduction by a superoxide reductase lacking the highly conserved lysine residue. J Biol Inorg Chem 2015; 20(1):155–64.

35. Horch M, Pinto AF, Utesch T, Mroginski MA, Romão CV, Teixeira M et al. Reductive activation and structural rearrangement in superoxide reductase: a combined infrared spectroscopic and computational study. Phys Chem Chem Phys 2014; 16(27):14220–30.

36. https://www.icds.si/wp-content/uploads/2018/09/P041_Troitzsch.pdf.

37. Warburg O, Christian W, Griese A. Hydrogen-transferring coenzyme, its composition and mode of action. Biochem Z 1935;282:157.

38. Braunersreuther V, Jaquet V. Reactive oxygen species in myocardial reperfusion injury: from physiopathology to therapeutic approaches. Curr Pharm Biotechnol 2012; 13(1):97–114.

39. Hulko M, Hospach I, Krasteva N, Nelles G. Cytochrome c biosensor--a model for gas sensing. Sensors (Basel) 2011; 11(6):5968–80.

40. Nauseef WM. Detection of superoxide anion and hydrogen peroxide production by cellular NADPH oxidases. Biochim Biophys Acta 2014; 1840(2):757–67.

41. Caulat LC, Lotoux A, Martins MC, Kint N, Anjou C, Teixeira M et al. Physiological role and complex regulation of O2-reducing enzymes in the obligate anaerobe *Clostridioides difficile*. mBio 2024:e0159124.

42. Anjou C, Lotoux A, Zhukova A, Royer M, Caulat LC, Capuzzo E et al. The multiplicity of thioredoxin systems meets the specific lifestyles of Clostridia. PLoS Pathog 2024; 20(2):e1012001.

43. Sebaihia M, Wren BW, Mullany P, Fairweather NF, Minton N, Stabler R et al. The multidrug-resistant human pathogen *Clostridium difficile* has a highly mobile, mosaic genome. Nat Genet 2006; 38(7):779–86.

44. Saujet L, Pereira FC, Serrano M, Soutourina O, Monot M, Shelyakin PV et al. Genome-wide analysis of cell type-specific gene transcription during spore formation in *Clostridium difficile*. PLoS Genet 2013; 9(10):e1003756.

45. Dembek M, Barquist L, Boinett CJ, Cain AK, Mayho M, Lawley TD et al. High-throughput analysis of gene essentiality and sporulation in *Clostridium difficile*. mBio 2015; 6(2):e02383.

46. Otto A, Maaß S, Lassek C, Becher D, Hecker M, Riedel K et al. The protein inventory of *Clostridium difficile* grown in complex and minimal medium. Proteomics Clin Appl 2016; 10(9-10):1068–72.

47. Mao GD, Poznansky MJ. Electron spin resonance study on the permeability of superoxide radicals in lipid bilayers and biological membranes. FEBS Lett 1992; 305(3):233–6.

48. Imlay JA. The molecular mechanisms and physiological consequences of oxidative stress: lessons from a model bacterium. Nat Rev Microbiol 2013; 11(7):443–54.

49. Ji C-J, Kim J-H, Won Y-B, Lee Y-E, Choi T-W, Ju S-Y et al. *Staphylococcus aureus* PerR Is a Hypersensitive Hydrogen Peroxide Sensor using Iron-mediated Histidine Oxidation. J Biol Chem 2015; 290(33):20374–86.

50. McQuade R, Roxas B, Viswanathan VK, Vedantam G. *Clostridium difficile* clinical isolates exhibit variable susceptibility and proteome alterations upon exposure to mammalian cationic antimicrobial peptides. Anaerobe 2012; 18(6):614–20.

51. Guedon E, Petitdemange H. Identification of the gene encoding NADH-rubredoxin oxidoreductase in *Clostridium acetobutylicum*. Biochem Biophys Res Commun 2001; 285(2):496–502.

52. Shetty N, Wren MWD, Coen PG. The role of glutamate dehydrogenase for the detection of *Clostridium difficile* in faecal samples: a meta-analysis. J Hosp Infect 2011; 77(1):1–6.

53. Girinathan BP, Braun SE, Govind R. *Clostridium difficile* glutamate dehydrogenase is a secreted enzyme that confers resistance to H2O2. Microbiology (Reading) 2014; 160(Pt 1):47–55.

54. Knippel RJ, Wexler AG, Miller JM, Beavers WN, Weiss A, Crécy-Lagard V de et al. *Clostridioides difficile* Senses and Hijacks Host Heme for Incorporation into an Oxidative Stress Defense System. Cell Host Microbe 2020; 28(3):411–421.e6.

55. Janoir C, Denève C, Bouttier S, Barbut F, Hoys S, Caleechum L et al. Adaptive strategies and pathogenesis of *Clostridium difficile* from in vivo transcriptomics. Infect Immun 2013; 81(10):3757–69.

56. Scaria J, Janvilisri T, Fubini S, Gleed RD, McDonough SP, Chang Y-F. *Clostridium difficile* transcriptome analysis using pig ligated loop model reveals modulation of pathways not modulated in vitro. J Infect Dis 2011; 203(11):1613–20.

57. Boyanova L, Boyanova L, Hadzhiyski P, Gergova R, Markovska R. Oxygen tolerance in anaerobes as a virulence factor and a health-beneficial property. Anaerobe 2024; 89:102897.

58. Dudek C-A, Jahn D. PRODORIC: state-of-the-art database of prokaryotic gene regulation. Nucleic Acids Res 2022; 50(D1):D295–D302.

59. Sievers F, Wilm A, Dineen D, Gibson TJ, Karplus K, Li W et al. Fast, scalable generation of high-quality protein multiple sequence alignments using Clustal Omega. Mol Syst Biol 2011; 7:539.

60. Naville M, Ghuillot-Gaudeffroy A, Marchais A, Gautheret D. ARNold: a web tool for the prediction of Rho-independent transcription terminators. RNA Biol 2011; 8(1):11–3.

61. Mirdita M, Schütze K, Moriwaki Y, Heo L, Ovchinnikov S, Steinegger M. ColabFold: making protein folding accessible to all. Nat Methods 2022; 19(6):679–82.

62. Castro E de, Sigrist CJA, Gattiker A, Bulliard V, Langendijk-Genevaux PS, Gasteiger E et al. ScanProsite: detection of PROSITE signature matches and ProRule-associated functional and structural residues in proteins. Nucleic Acids Res 2006; 34(Web Server issue):W362-5.

63. Thoma S, Schobert M. An improved *Escherichia coli* donor strain for diparental mating. FEMS Microbiol Lett 2009; 294(2):127–32.

64. Ransom EM, Ellermeier CD, Weiss DS. Use of mCherry Red fluorescent protein for studies of protein localization and gene expression in *Clostridium difficile*. Appl Environ Microbiol 2015; 81(5):1652–60.

65. Kruger NJ. The Bradford method for protein quantitation. Methods Mol Biol 1994; 32:9–15.

66. Sievers S, Dittmann S, Jordt T, Otto A, Hochgräfe F, Riedel K. Comprehensive Redox Profiling of the Thiol Proteome of *Clostridium difficile*. Mol Cell Proteomics 2018; 17(5):1035–46.

67. McCord JM, Fridovich I. The utility of superoxide dismutase in studying free radical reactions. II. The mechanism of the mediation of cytochrome c reduction by a variety of electron carriers. J Biol Chem 1970; 245(6):1374–7.

