## Supplementary material for "A rubrerythrin locus of *Clostridioides difficile* efficiently detoxifies reactive oxygen species": Knop et al_supplements

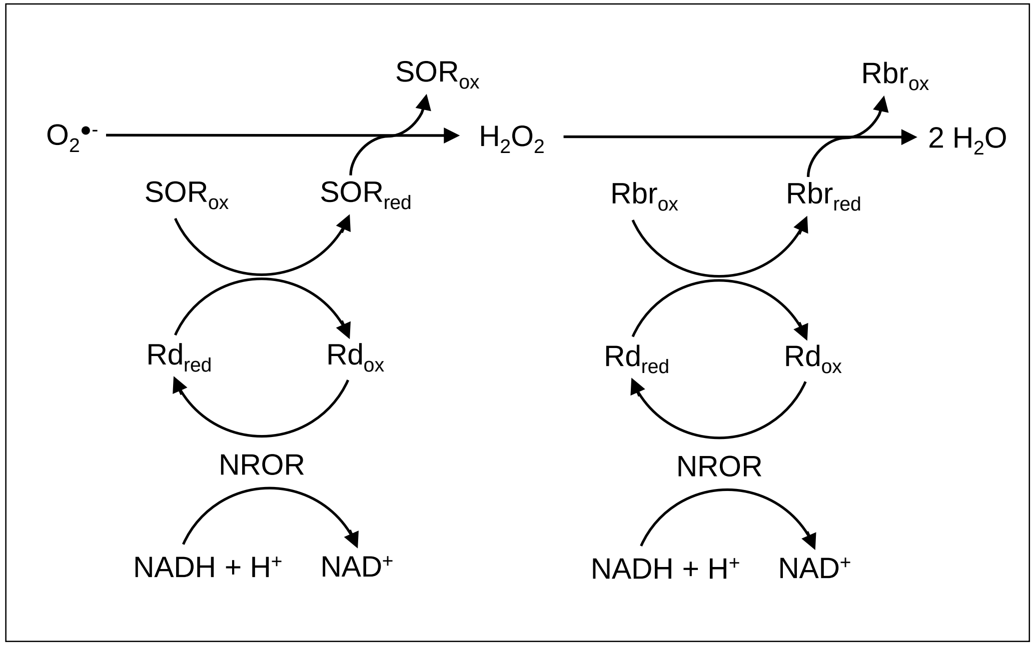


**Figure S1:** Reductive detoxification of $O_2^{\bullet -}$ and reactive $H_2O_2$. NROR: NADH:Rubredoxine Oxidoreductase; Rbr: Ruberythrin; Rd: Rubredoxin; SOR: Superoxid reductase; red: reduced; ox: oxidized. (modified from Hillmann 2009, (18)).

```
                                   Fe Center I
C. difficile 630   ----MCSEQKFFICETCGNLVGMIQSGGVPIFCCGKPMKELVPNTTDAAVEKHVPVIEVD    56
C. acetobutylicum  ----MNNDLSIYVSKNSGTAVLLLQGNGTDLTCGSEPMAKIVANTTDAAQEKHVPHITKN    56
C. beijerinckii    MMKGGYIMIAFYRCELCGNIVGLIKNGGGQLVCCGKNMTKLEANSTDAAQEKHVPVAERK    60
C. tetani          ----MTELLQVYKCQVCGNIVEVVHKGGGQLVCCNQPMKLFVENTVDAAVEKHVPVIEKI    56
C. botulinum       ----MVKLNEVYKCEVCGNIVQVVHASGGQLVCCGKPMRLLEENTTDAALEKHVPVVEKT    56
                             .: .: .*. * ::: .*  : * .: *  :  *:.*** *****

                                            Fe Center II
C. difficile 630   GNNVTVKVSSTTHPMTKEHHIAWVYLMTEQGGQRKCLAVDGEPVVKFALNDDDKVISAYA    116
C. acetobutylicum  GNNIDVSVGSVEHPMTPEHFIEWIILVSGDRLEMAKLTPDMKPRAQFHN---VTSGTVYA    113
C. beijerinckii    DGKIYVTVGSVEHPMTEEHYIEWIAVVSDKGTEKVSLSPGEKPQAVFVD---KGNAVVYA    117
C. tetani          EGGIRVKIGEAEHPMIEEHYIEWIEVLTENKVYRKHLKPGEKPVAEFKL--DEEVVAARE    114
C. botulinum       DNGIKVKVGEKEHPMEEKHYIEWIEVITEDKVYKKYLKPGEKPEAEFKL--DEEVVKVRE    114
                    . : *.:..  ***  :*.* *: ::: .      *  . :* . *          .

C. difficile 630   YCNLHGLWKAEL       128
C. acetobutylicum  YCNLHSLWKADI       125
C. beijerinckii    YCNLHGLWKTE-       128
C. tetani          YCNLHGLWKK--       124
C. botulinum       YCNIHGLWKK--       124
                   ***:*.***
```

**Figure S2**: Amino acid alignment of *C. difficile* 630 Rbo to homologs of other clostridia. Iron binding amino acids are colored in yellow for the rubredoxin-type Fe center I [Fe(SCys)$_4$] and turquoise for the Fe center II [Fe(NHis)$_4$(SCys)]. asterisk: completely similar; colon: very similar; dot: similar

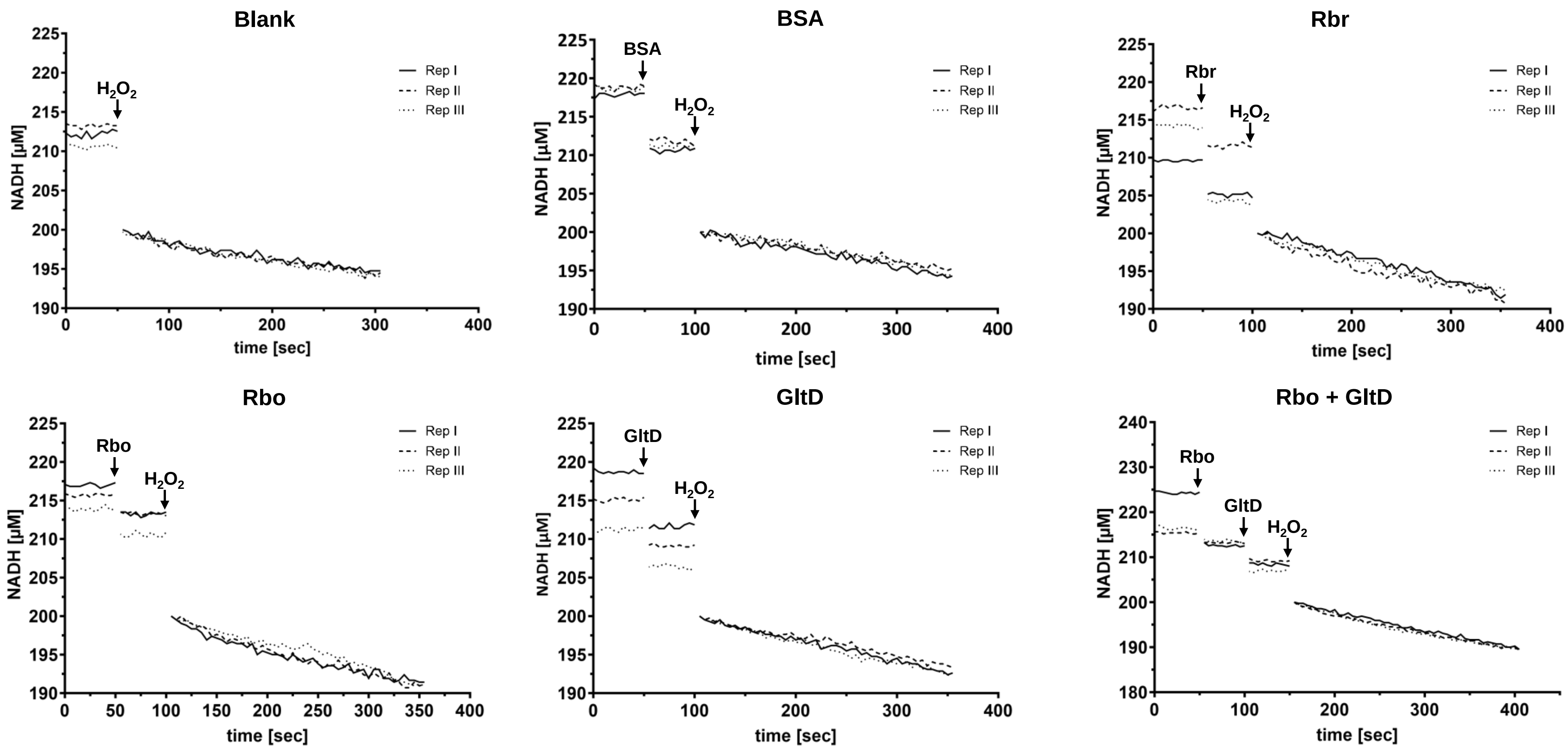


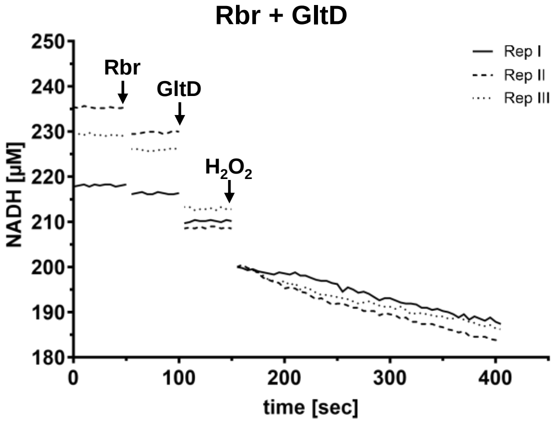


**Figure S3:** $H_2O_2$ reductase activity of BSA, Rbr, Rbo, GltD. The activities were determined anaerobically by measuring NADH consumption monitored at 340 nm every 5 seconds. Each assay was performed in three replicates. Shown are BSA, Rbr, Rbo and GltD individually, as well as the measured values of the combinations of Rbr + GltD and Rbo + GltD. The protein concentrations in the assays were 1 µM for each enzyme (BSA, Rbr, Rbo, GltD). For the blank experiment only NADH and $H_2O_2$ in buffer C were in the cuvette. Experiments were performed in the presence of 200 µM NADH and 100 µM $H_2O_2$. Before the addition of $H_2O_2$,10 measurements with each compound were taken to see a constant NADH level and after addition normalized to 200 µM NADH.

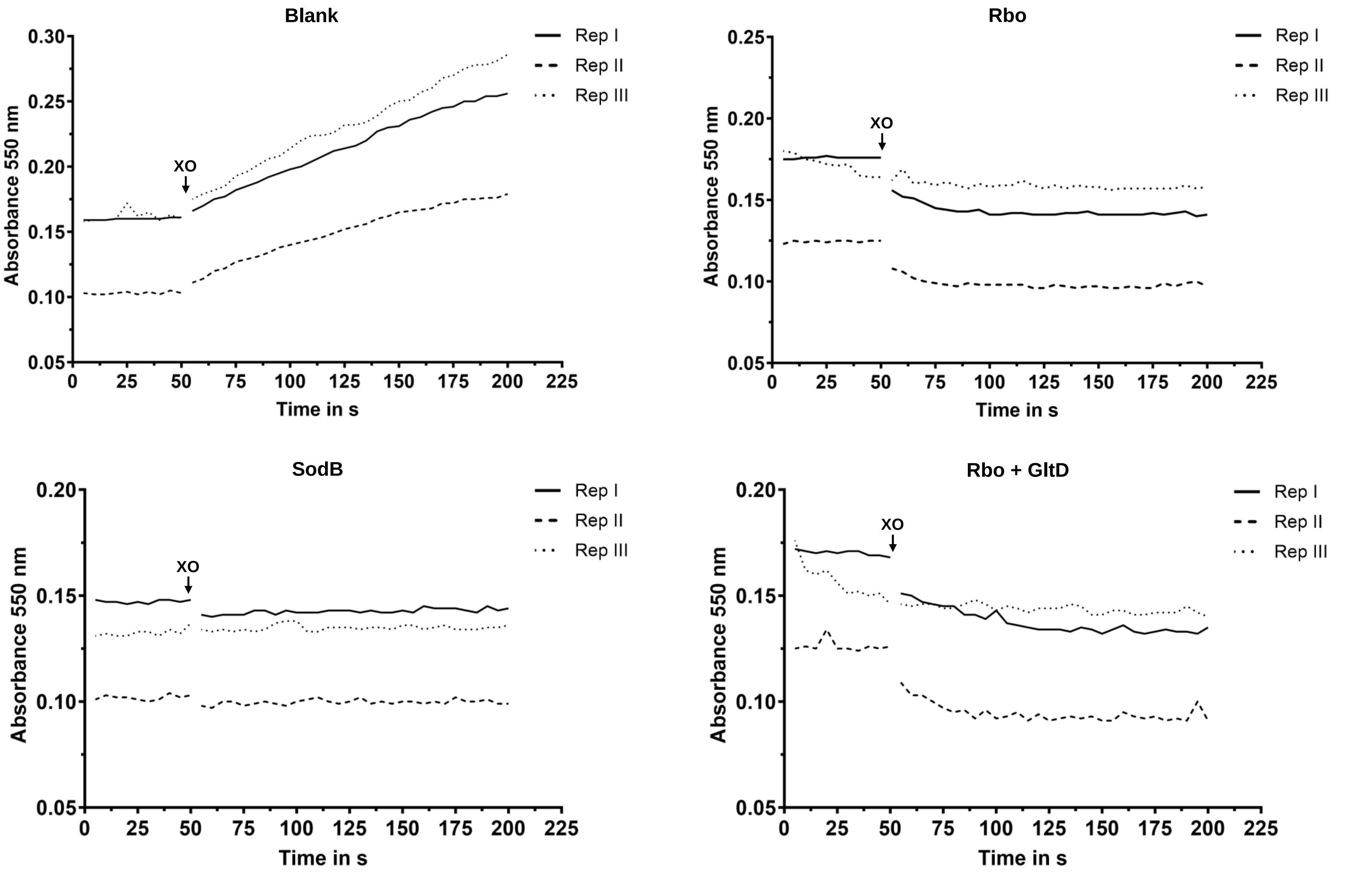


**Figure S4:** Superoxide reductase activity of Rbo, SodB and Rbo + GltD measured via absorbance at 550 nm every 5 seconds indicating cytochrome c reduction. Shown are Rbo and SodB individually, as well as the measured values of the combination of Rbo + GltD. The protein concentrations in the assays were 1 µM for each enzyme (Rbo, GltD, SodB). Experiments were performed in the presence of 200 µM NADH, 500 µM xanthine, 20 µM cytochrome c and 15 µg/ml xanthine oxidase (XO) added to start the generation of superoxide. Each assay was performed in three replicates.

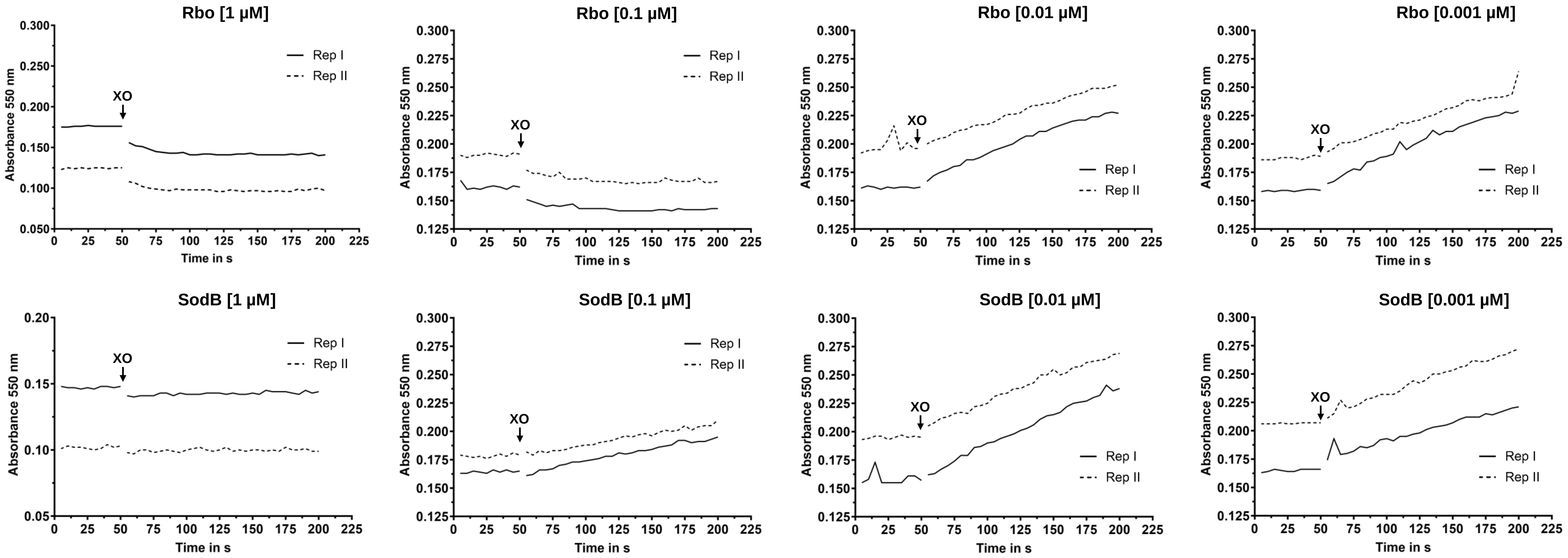


**Figure. S5:** Superoxide reductase activity of Rbo and SodB measured via absorbance at 550 nm every 5 seconds in a concentration gradient. Protein concentrations of 1 µM; 0.1 µM; 0.01 µM and 0.001 µM were tested. Each assay was performed in two replicates. Experiments were carried out in the presence of 200 µM NADH, 20 µM cytochrome c, 500 µM xanthine and 15 µg/ml xanthine oxidase (XO) for the superoxide generation.

**Table S1**: Blast analysis of GltD homologs of other species than *C. difficile*. Shown are 20 species with the highest percent identity (status 20th of June 2024).

| Description | Species | % identity to *C. difficile* 630 | E Value | Bit score | Query cover [%] | Acc. Len | Accession |
|---|---|---|---|---|---|---|---|
| Glutamate synthase-related protein | *Clostridium argentinense* | 76.67 | 0 | 773 | 100 | 470 | WP_039633109.1 |
| Glutamate synthase-related protein | *Metaclostridioides mangenotii* | 71.55 | 0 | 730 | 99 | 475 | WP_216467922.1 |
| Glutamate synthase-related protein | *Anaerofustis stercorihominis* | 68.68 | 0 | 659 | 99 | 472 | WP_306813826.1 |
| Glutamate synthase-related protein | *Zongyangia hominis* | 67.78 | 0 | 649 | 100 | 470 | WP_262396883.1 |
| Glutamate synthase-related protein | *Faecalicatena faecalis* | 67.57 | 0 | 652 | 99 | 468 | WP_216243410.1 |
| Glutamate synthase-related protein | *Gottschalkia acidurici* | 67.43 | 0 | 659 | 99 | 467 | WP_014966750.1 |
| Glutamate synthase-related protein | *Diplocloster hominis* | 67.22 | 0 | 669 | 99 | 461 | WP_343249707.1 |
| Glutamate synthase-related protein | *Robinsoniella peoriensis* | 67.08 | 0 | 673 | 99 | 480 | WP_243132993.1 |
| Glutamate synthase-related protein | *Diplocloster agilis* | 67.01 | 0 | 674 | 99 | 461 | WP_238721002.1 |
| Glutamate synthase-related protein | *Suonthocola fibrivorans* | 67.01 | 0 | 673 | 99 | 461 | WP_158347964.1 |
| Glutamate synthase-related protein | *Diplocloster modestus* | 66.81 | 0 | 667 | 99 | 461 | WP_158352183.1 |
| Glutamate synthase-related protein | *Anaerotignum faecicola* | 66.81 | 0 | 654 | 99 | 466 | MCQ4726245.1 |
| Glutamate synthase-related protein | *Monoglobus pectinilyticus* | 66.81 | 0 | 653 | 99 | 475 | WP_102366369.1 |
| Glutamate synthase-related protein | *Methanobrevibacter ruminantium* | 66.67 | 0 | 654 | 100 | 470 | WP_012955612.1 |
| Glutamate synthase-related protein | *Qiania dongpingensis* | 65.97 | 0 | 659 | 99 | 464 | WP_249301298.1 |
| Glutamate synthase-related protein | *Lactonifactor longoviformis* | 65.62 | 0 | 656 | 100 | 463 | WP_072854469.1 |
| Glutamate synthase-related protein | *Hungatella effluvii* | 65.42 | 0 | 657 | 99 | 476 | WP_110321870.1 |
| Glutamate synthase-related protein | *Hungatella hathewayi* | 65.42 | 0 | 657 | 99 | 476 | WP_002604813.1 |
| Glutamate synthase-related protein | *Hungatella hominis* | 65.00 | 0 | 654 | 99 | 476 | WP_187020201.1 |
| Glutamate synthase-related protein | *Lientehia hominis* | 64.93 | 0 | 657 | 99 | 464 | WP_231062982.1 |

**Table S2:** Oligonucleotides used in this study.

| Name | Sequence | Function |
|---|---|---|
| rbr1+RBS_pDSW1728_SacI_F | ATGGTAGAGCTCAATATAATGTTGGGAGGAATTTAAGAAATGAACTTAAAAGGAACTAAAACAGAA | construction of *rbr*-pDSW1728 overexpression plasmid |
| Rbr1_Twin-Streptag_Fusion_R | CTGCGGGTGGCTCCAAGCGCTCCCATAATTTTCAGCTTTTATATTAAAATATGC | |
| Twin-Streptag_Rbr1_Fusion_F | GCATATTTTAATATAAAAGCTGAAAATTATGGGAGCGCTTGGAGCC | |
| Twin-Streptag_pDSW1728_BamHI_R | ATGGTAGGATCCTTATTTCTCGAACTGCGGGTGG | |
| dfx+RBS_pDSW1728_SacI_F | ATGGTAGAGCTCAATATAATGTTGGGAGGAATTTAAGAAATGTGTAGTGAACAAAAATTTTTTATATGTG | construction of *dfx*-pDSW1728 overexpression plasmid |
| dfx_Twin-Streptag_Fusion_R | CTGCGGGTGGCTCCAAGCGCTCCCCAGTTCAGCCTTCCATAATCCATG | |
| Twin-Streptag_dfx_Fusion_F | TATCAAATTATACAAACATACAGCATGTAGGGAGCGCTTGGAGCC | |
| CD630_08280+RBS_pDSW1728_SacI_F | ATGGTAGAGCTCAATATAATGTTGGGAGGAATTTAAGAAATGTCAATTTATAAATGTAGTGTTTGTGG | construction of *CD630_08280*-pDSW1728 overexpression plasmid |
| CD630_08280_Twin-Streptag_Fusion_R | CTGCGGGTGGCTCCAAGCGCTCCCTACATGCTGTATGTTTGTATAATTTGATA | |
| Twin-Streptag_CD630_08280_Fusion_F | TATCAAATTATACAAACATACAGCATGTAGGGAGCGCTTGGAGCC | |
| EC_sodB_Esp3I_For | GGAGATATACAATGTCATTCGAATTAC | construction of *sodB*-pIBA103 overexpression plasmid |
| EC_sodB_Esp3I_Rev | CTCCAAGCGCTCCCTGCAGCGAGATTT | |
| IBA103_seq_For | GAGTTATTTTACCACTCCCT | sequence control of pIBA103 |
| IBA103_seq_Rev | CGCAGTAGCGGTAAACG | |
| pDSW1728_control_F | GCTTGATCGTAGCGTTAACAG | sequence control of pDSW1728 |
| pDSW1728_control_R | CTTCTCATGAGAGAAGCCTTTTTC | |

**Table S3:** Plasmids used in this study.

| Name | Discription | Origin |
|---|---|---|
| pDSW1728 | *E. coli-C. difficile* shuttle vector: tetracycline-inducible promoter (Ptet) driving expression of red fluorescent protein (mCherryOpt) | doi: 10.1128/AEM.03446-14 (64) |
| pASG-IBA103 | Cytoplasmic *E. coli* expression vector encoding a C-terminal Twin-Strep-tag® | IBA Lifesciences GmbH |
| *rbr*-pDSW1728 | *E. coli-C. difficile* shuttle vector: tetracycline-inducible promoter (Ptet) driving expression of *rbr* | in this study |
| *dfx*-pDSW1728 | *E. coli-C. difficile* shuttle vector: tetracycline-inducible promoter (Ptet) driving expression of *dfx* | in this study |
| *CD630_08280*-pDSW1728 | *E. coli-C. difficile* shuttle vector: tetracycline-inducible promoter (Ptet) driving expression of *CD630_08280* | in this study |
